# Carbon concentrating mechanisms in a small coastal diatom demonstrate sensitivity to changes in Zinc and carbonate chemistry

**DOI:** 10.64898/2026.05.21.726966

**Authors:** Emily A. Burdige, Mak A. Saito, Matthew G. Hayden, Paloma Z. Lopez, Matthew R. McIlvin, Adam V. Subhas

## Abstract

The efficiency of marine diatom carbon concentrating mechanisms (CCMs) play a critical role in photosynthesis, enabling cells to maintain rapid growth rates under a variety of environmental conditions. To assess the plasticity of the model diatom *P. tricornutum*’s CCMs, specifically carbonic anhydrase (CA) enzymes and bicarbonate transporters, we measured growth response, bulk CA activity, and corresponding shifts in the proteome under a range of Zn and pCO_2_ conditions in trace metal-controlled culture. CA activity increased with Zn availability and decreased with pCO_2_. A positive growth effect was observed due to Zn addition and increasing pCO_2_ from 200 to 400 ppm, however growth rate decreased as pCO_2_ further increased to 750 ppm. Across the six treatments, the protein abundance of ISIP2A, which functions to bring inorganic Fe into the cell via a FeCO_3_ complex and is used as a biomarker for Fe stress, demonstrated an inverse relationship with [CO_3_^2-^], consistent with its role as a phytotransferrin. Under conditions of Zn limitation ([Zn^2+^] = 0.3 pM), the cell appeared to allocate this metal away from CA, instead relying on a uniquely expressed CA isoform with a 100-fold lower intrinsic activity than that of the primary Zn-CA, as calculated using paired abundance-activity measurements. We furthermore observed a continued increase in bicarbonate transport protein abundance after CA activity plateaued, suggesting any deficit in DIC required to maintain growth rates was accomplished through HCO_3_^-^ transport. We hypothesize bicarbonate uptake and CO_2_ diffusion operate in tandem via CA enzymatic activity to supply adequate CO_2_ for photosynthesis.

## Introduction

Diatoms are a prolific group of photoautotrophic algae that contribute to upwards of 40% of marine primary production (Tréguer et al. 2018), exerting a strong lever on surface water pCO_2_ (Cai et al. 2020). During bloom events, phytoplankton rapidly fix aqueous carbon dioxide (CO_2_), into biomass typically faster than the pool of dissolved inorganic carbon (DIC) is able to re-equilibrate (Tortell et al. 2011). Moreover, due to their dense Si frustule, aggregates of diatom-associated organic matter rapidly sink out of the surface ocean, providing an efficient mechanism for CO_2_ export to the deep ocean (Smetacek 1999). Together, these processes give diatoms a critical role in the overall biological carbon pump.

At high growth rates, phytoplankton must maintain a sufficient flux of CO_2_ from seawater to the site of C fixation in the chloroplast (Badger and Price 1994). This intracellular carbon transport across multiple membranes is facilitated by carbonic anhydrase (CA) metalloenzymes, that are one component of cells’ carbon concentrating mechanisms (CCMs) and operate to catalyze the equilibration of the DIC pool (see below). Thus far, there are eight identified subclasses of CAs (*α*, *β*, *γ*, *δ*, *ζ*, *η*, *θ*, and *ι*), seven of which exist in diatom genomes (*α*, *β*, *γ*, *δ*, *θ*, *ζ,* and *ι*), all of which perform the same function, but share little sequence or structural similarity to each other (DiMario et al. 2018a; Jensen et al. 2019). Zinc (Zn) is the preferred and canonical cofactor for CA, making it an essential micronutrient for diatom photosynthesis; however, some diatoms possess the ability to substitute different divalent metals in the CA active site to maintain activity. The diatom *Thalassiosira weissflogii* possess two CAs which can replace Cd, (ζ-CA, CDCA) or Co (δ-CA, TWCA1) for Zn (Price and Morel 1990; Yee and Morel 1996). Similarly, an *ι*-CA was identified in *T. pseudonana* which unusually has a higher activity with Mn in its active site relative to Zn^2+^, Ca^2+^, Cd^2+^, Co^2+^, or Mg^2+^ (Jensen et al. 2019). Recently, the abundance of *ι*-CA in *P. tricornutum* was identified to increase under low Zn and pCO_2_ conditions, but operate without a metal cofactor (Avilan et al. 2026). Though there now exists conflicting evidence about the role of metals in CA, the prevailing hypothesis is that when those replacement elements are similarly scarce, growth becomes limited (Kellogg et al. 2022; Morel et al. 1994; Saito et al. 2008).

The mechanisms for inorganic carbon uptake and utilization by marine phytoplankton remain a dynamic area of research. Seawater DIC concentrations in the form of bicarbonate ion (HCO_3_^-^) are 200-fold higher than that of CO_2_(aq), raising the question of whether phytoplankton can actively take up HCO_3_^-^ followed by internal conversion to CO_2_ for photosynthesis. The uncatalyzed hydration of CO_2_ takes 28 seconds at 25°C and salinity of 34 (Johnson 1982)

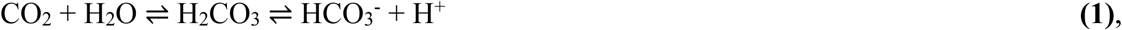

and is kinetically slow. However, the product, H_2_CO_3_, dissociates essentially instantaneously, as the acid-base reactions that equilibrate H_2_CO_3_, HCO_3_^-^, and CO_3_^2-^ proceed via rapid (de)-protonation. If the intracellular flux of DIC is limited to CO_2_ diffusion, its rate of consumption by RuBisCO in nutrient-replete conditions can exceed the rate at which HCO_3_^-^ abiotically equilibrates with CO_2_, resulting in a kinetic limitation on phytoplankton growth (Riebesell et al. 1993). Further, the K_M_ of CO_2_ for RuBisCO is approximately 40 μM, which is 4-times higher than the average concentration of CO_2_(aq) in surface seawater (Young et al. 2016). To adapt to both the kinetic and thermodynamic constraints on CO_2_ uptake and utilization, cells have evolved highly efficient CCMs that operate to accumulate CO_2_ in the vicinity of RuBisCO and are the linchpin of C uptake in marine phytoplankton. These CCMs include multiple CAs that catalyze the equilibration of CO_2(aq)_ and H_2_CO_3_, helping to alleviate kinetic limitations on DIC uptake from seawater and utilization within the cell. These CAs require metals to function, providing a mechanistic linkage between CO_2_ and metal limitation in marine phytoplankton (Saito et al., 2008).

CAs are located within the cell among various organelles and sometimes are expressed extracellularly (eCA) at the cell membrane where they can interact with the seawater in the diffusive boundary layer (DBL) surrounding the cell (Keys et al. 2024). As CO_2_ is rapidly depleted in the seawater within the DBL, eCA catalyzes its equilibration with HCO_3_^-^, which contributes to the concentration gradient that enables the diffusive flux into the cell. Within the cell, intracellular CAs (iCAs) work to funnel CO_2_ through the cytosol and to the chloroplast (Badger and Price 1994; Satoh et al. 2001). A feature among certain algae with CCMs, including *Phaeodactylum tricornutum*, is the pyrenoid, a subcompartment of the chloroplast encased in a proteinaceous sheath that functions to further concentrate CO_2_ in the vicinity of RuBisCO enzymes (Shimakawa et al. 2024). In contrast to CAs, the pyrenoid is not ubiquitous among all algae with CCM activity (DiMario et al. 2018b). An additional component of phytoplankton cell’s CCM are specific solute carrier proteins for the direct uptake of HCO_3_^-^, where it is subsequently converted to CO_2_ prior to fixation by RuBisCO (Nakajima et al. 2013; Tortell et al. 1997). Discriminating between CA-mediated CO_2_ diffusion and HCO_3_^-^ uptake remains a challenge, as the balance between the two mechanisms varies among species (e.g., Burkhardt et al. 2001; Martin and Tortell 2008; Rost et al. 2003; Trimborn et al. 2009).

The coastal diatom *Phaeodactylum tricornutum* is known to maintain rapid growth rates and reach high cell densities in nutrient-replete conditions, making it an ideal organism to study under conditions of CO_2_ drawdown. Moreover, its genome is fully sequenced (Bowler et al. 2008), and there is a robust field of literature which aims to understand its physiology. With regards to C acquisition strategies in other diatoms, prior studies have demonstrated increases in CA activity and HCO_3_^-^ uptake at low pCO_2_ (Burkhardt et al. 2001; Rost et al. 2003; Trimborn et al. 2009) and increases in CA protein expression in replete Zn concentrations (Kellogg et al. 2022; W. Li et al. 2020). Fewer studies have simultaneously manipulated Zn concentration and pCO_2_ (e.g., Lane & Morel, 2000; Morel et al., 1994; Sunda & Huntsman, 2005), none of which have been carried out on *P. tricornutum* to our knowledge. Within the multiple strains of *P. tricornutum*, there is a variable proportion of extracellular CA activity relative to total enzyme activity; in ten of the eleven strains studied, this proportion was under 20% (John-McKay and Colman 1997). Additionally, genomic work has identified one strain of *P. tricornutum* lacks genes for any external CAs (Samukawa et al. 2014; Tachibana et al. 2011). There appears to be a general agreement that this diatom relies heavily on intracellular CAs and the uptake of HCO_3_^-^, providing a unique perspective for studying cellular CCMs (Satoh et al. 2001; Nawaly et al. 2023). This study examines the influence of Zn and pCO_2_ on *P. tricornutum’s* CCMs, specifically changes to its proteome and resultant CA enzyme activity.

## Methods

Two separate culturing experiments were conducted with the diatom *P. tricornutum*. In a CO_2_ manipulation experiment, cultures were grown at three pCO_2_ conditions (200 ppm CO_2_, 400 ppm CO_2_, 750 ppm CO_2_, henceforth low, mid and high CO_2_; chosen in accordance with the *Guide to Best Practices for Ocean Acidification Research* (Riebesell et al. 2011)), and amended with 100 nM total dissolved Zn ([Zn]_T_). See below for details on the ebullition design for these experiments. In a Zn manipulation experiment, cultures were grown at three different total Zn concentrations (3 nM, 30 nM, and 100 nM [Zn]_T_, henceforth low, mid and high Zn) without any seawater pCO_2_ modification (e.g. via bubbling), but with bottle caps loosely threaded to allow gas exchange with ambient air. In both experiments, cell count measurements were made daily via flow cytometry and samples were taken for Total Alkalinity (TA) and DIC immediately prior to harvesting via centrifugation for proteomics and carbonic anhydrase activity during late-log phase.

### Culturing

All culture work was carried out in a Class 100 trace metal clean room. Polycarbonate bottles were trace metal cleaned (4-day soak in <1% Citranox detergent, 5 rinses in Milli-Q water, a 7-day soak in 10% HCl, 2 rinses with dilute acid (HCl, pH 2), and >5 rinses with Milli-Q) and microwave sterilized prior to use. All solutions were added after rinsing the pipette tips to remove any trace metals (this consisted of 1 rinse of 10% HCl, 3 rinses with sterile dilute HCl (pH 2), and >5 rinses with Milli-Q water).

A modified f/2 media base was used for all culturing. The base of the media was 0.2 µm-filtered and microwave sterilized seawater collected from the North Pacific Ocean at St. 17 (25.082° N, 118.101° W; salinity 33.9) on the CliOMZ cruise (Lopez et al. 2024). Macronutrients were added to the culture bottles to reach final concentrations of 88.2 µM NaNO_3_, 36.2 µM NaH_2_PO_4_, and 106 µM Na_2_SiO_3_. The vitamin composition consisted of 2 nM biotin, 0.37 nM B_12_ as cyanocobalamin, and 300 nM thiamine. The vitamin and macronutrient solutions were sterile filtered and chelexed prior to addition to seawater. Finally, a sterile filtered trace metal mix was added to the media with final concentrations of 4.8×10^-8^ M MnCl_2_, 10^-8^ M Na_2_O_3_Se, 4.0×10^-8^ M CuSO_4_, 10^-7^ M FeCl_3_, and 10^-7^ M NiCl_2_, in a 10^-4^ M ethylenediamine tetraacetic acid disodium salt (EDTA, Acros Organics, HV-88113-90) metal ion buffer system. The trace metal mix used for culture work was made without the addition of Zn or Co such that the Zn concentration could be manipulated on an experimental basis, and without substitution by Co. The modified f/2 media base was amended with total added Zn concentrations of 3, 30, and 100 nM. Total Zn is a summation of all Zn species, including those bound by organic and inorganic ligands and free Zn^+2^ ions. The ratio of free Zn^2+^ to the total dissolved Zn in seawater are proportional to one another and equal 10^−3.99^ as used by Sunda and Huntsman (1995). The corresponding [Zn^2+^] values are 0.3, 3, and 10 pM. EDTA in the seawater media prevents large changes in [Zn^2+^] due to pH and throughout the growth period as biomass increases (see Supporting Information Fig. S1, S2, and Table S1).

**Fig. 1.**
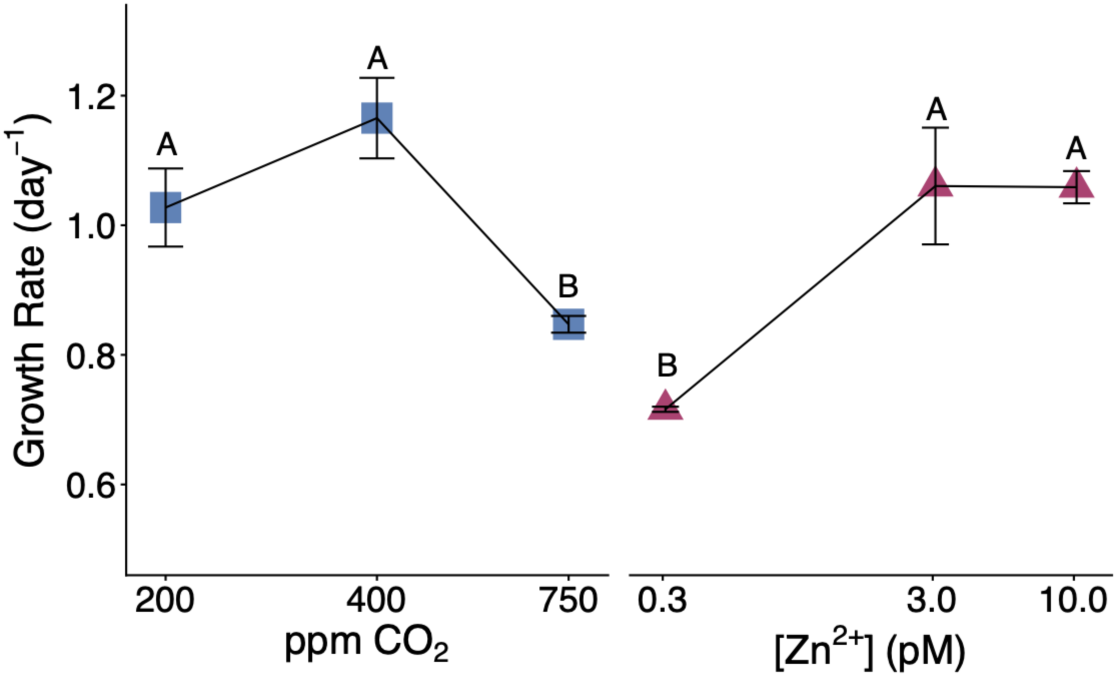
Growth rates across the six treatments with pairwise comparison significance (ANOVA *p* = 2.2E-6). Data are reported as the average and standard deviation of biological triplicate. Treatments with matching letters reflect an insignificant difference in protein abundance from one another (*p* < 0.05; ANOVA followed by Tukey post hoc test).

**Fig. 2.**
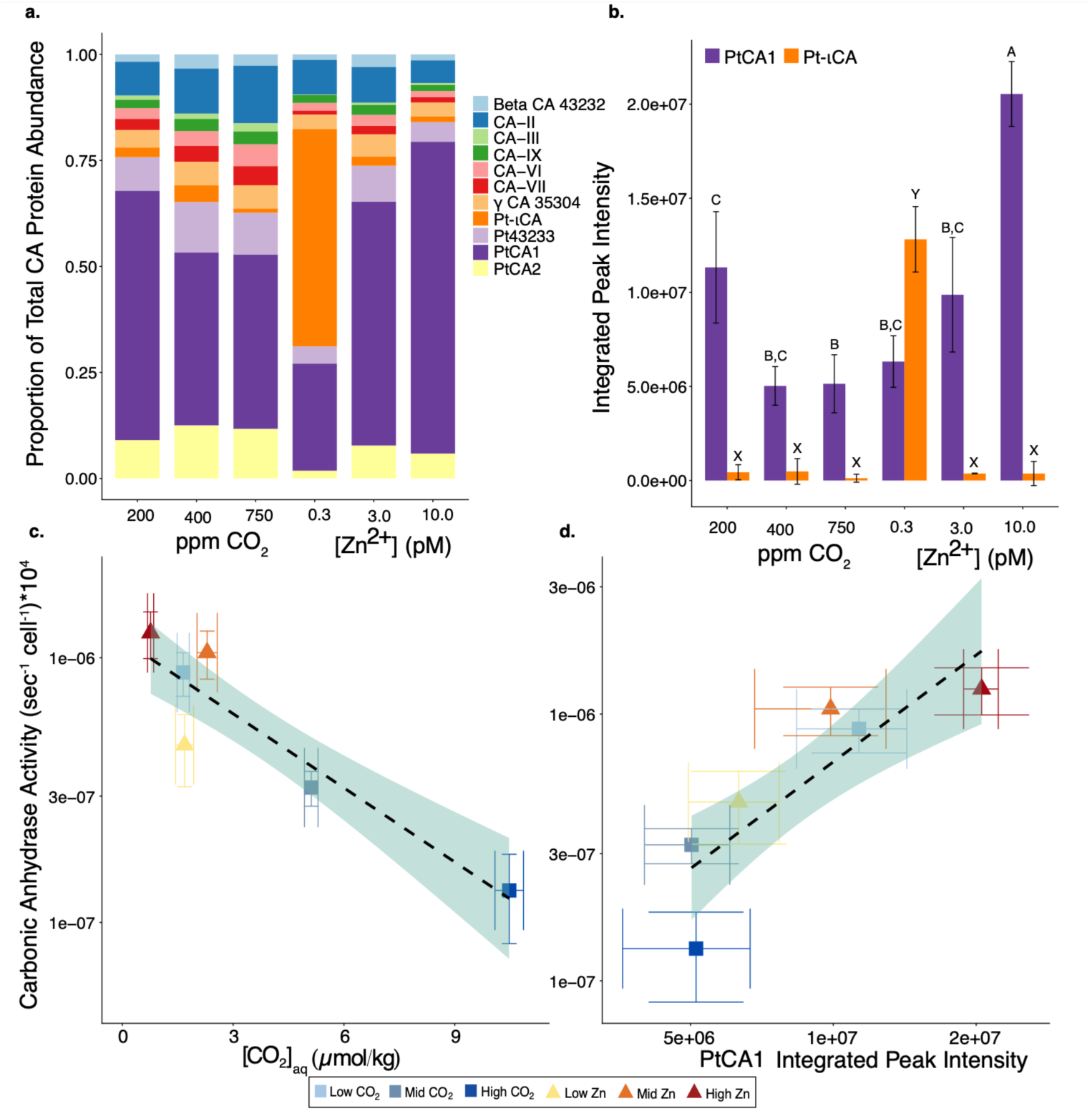
Carbonic Anhydrase protein abundances and activity response to treatments. (**a**) Normalized abundances of all identified CA proteins for each experiment. (**b**) Protein abundance of PtCA1 (purple) and Pt-ιCA (orange). Treatments with differing letters (A/B/C for PtCA1; X/Y for Pt-ιCA) reflect significant differences from one another (*p* < 0.05; ANOVA followed by Tukey post hoc test). (**c**) CA activity plotted against the corresponding CO_2_(aq) concentration in the seawater and (**d**) mean protein abundance of PtCA1. Dashed line represents a linear regression model with an 80% confidence band in light green. Data are reported as the average and standard deviation of biological triplicate (standard deviation omitted in panel a). For further information regarding regression parameters, refer to Supporting Information Table S6.

*Phaeodactylum tricornutum* CCMP632 was obtained from the National Center for Marine Algae and Microbiota at Bigelow Laboratory for Ocean Sciences. All cultures were grown in a 25.5°C incubator with 24 hr day^-1^ fluorescent lighting. Neutral density mesh screening was used to filter the light source to 80 μmol photon m^−2^ s^−1^. Stock cultures at the aforementioned Zn concentrations were maintained without bubbling in trace metal-cleaned, loosely capped 28 mL polycarbonate tubes prior to experimental work and were used as inoculum for all experiments. For the two experiments, cultures were grown in 250 mL polycarbonate bottles to accommodate the volume required to harvest cells for proteomics and CA activity. Biomass was harvested via centrifugation (3,050 × g for 30 min at 4°C in 50 mL centrifuge tubes followed by decantation of the supernatant and an additional centrifugation at 14,100 × g for 8 min at room temperature in 2 mL centrifuge tubes) and immediately frozen at – 80°C where they were stored until processing to prevent protein degradation and subsequent loss of CA activity. Separate cell pellets were collected for measurement of CA activity and protein extraction.

To monitor growth, the cell densities of all cultures were measured daily using a flow cytometer (BD Accuri C6 Plus) and gated according to their Forward Scatter and PerCP-A (Peridinin-Chlorophyll-Protein A) concentration. At cell densities exceeding 10^4^ cells mL^-1^, the aliquot taken for flow cytometry was diluted down for an accurate cell concentration measurement.

Aerated experimental treatments were bubbled with 400 mL min^-1^ of CO_2_-free air, as determined by an abiotic evaporation experiment (Supporting Information Fig. S3) and mixed with different ratios of 1% (10,000 ppm) CO_2_ to reach desired pCO_2_ values of 200, 400, 750 ppm. Prior to any experiments, the pCO_2_ of the gas stream that was delivered to the seawater was measured and validated using a LICOR (LI-7815) to ensure it was equivalent to the treatment pCO_2_ set point. A schematic of the aeration apparatus can be found in Supporting Information Fig. S4.

Seawater samples were taken for TA and DIC at the termination of the experiment to examine the carbonate chemistry profile of the seawater at the same time point at which samples were taken for CA activity and proteomics. Further methods on sample analysis for DIC and TA can be found in Supporting Information Text S1.

### Carbonic Anhydrase Activity Measurements via Membrane Inlet Mass Spectrometer

Biomass pellets for measurements of CA activity contained at least 10^6^ cells and were analyzed within one week of collection. All samples were assayed for their CA activity following a ^18^O-depletion method was followed, which tracks the ion currents of three different ^13^C and ^18^O labeled CO_2_ isotopes over time as a double-labeled spike solution (NaH^13^C^18^O_3_, m/z 49) interacts with the background buffer solution. Measurements were made on a membrane inlet mass spectrometer (MIMS), which measures ion intensities of dissolved gases based on detection of their mass to charge (m/z) ratio (Burlacot et al. 2020). Further details about the CA activity method can be found in Supporting Information Text S2.

Bovine carbonic anhydrase (BCA) was used to create a calibration between known concentrations of BCA and the measured rates of CA activity. Standard curves were used to assess instrument performance but were not used to calculate an effective CA concentration for samples, as the activity of a known concentration of bovine carbonic anhydrase is incomparable to phytoplankton CAs (Krishnamurthy et al. 2008). Additionally, “blank” measurements of buffer containing no standard addition of BCA were run every analytical session. The mean plus 1σ standard deviation of all the blanks provided a conservative estimate of the variability in these measurements. This value was subtracted from the sample’s measured CA activity to achieve the blank subtracted CA activity. The data in this manuscript are reported as blank subtracted CA activity multiplied by 10^4^ (to achieve a whole number) and normalized to the number of cells analyzed for the respective sample, with units of reactions sec^-1^ cell^-1^.

### Proteomic Analysis

Harvested cell pellets were removed from the freezer and resuspended in lysis buffer for protein extraction. The total extracted protein was quantified, and 25 μg were subsampled for the reduction, alkylation, and digestion of proteins. Subsequently, samples were transferred to an S-trap column (Protifi, S-Trap Mini Column) to clean, isolate, digest, and elute the proteins for analysis. Tryptic peptides were analyzed with liquid chromatography coupled with tandem mass spectrometry (LS/MS/MS) and data independent acquisition (DIA) was used for analysis of the biological triplicates. For detailed methods of all proteomic analysis, refer to Supporting Information Text S3.

### Proteomic Data Processing

Protein abundances are reported as the integrated peak intensity for peptides associated with each protein averaged between biological triplicates (Maximum Label-Free Quantification in DIA-NN). In instances where summed protein abundances are reported (i.e. for CA or bicarbonate transport proteins), the average abundances were added together for each treatment, and the error was propagated accordingly. The mean number of uniquely identified proteins across all biological replicates for each treatment was 5961. One technical replicate in the Mid CO_2_ treatment identified significantly fewer unique proteins relative to all other samples, due to an insufficient protein concentration analyzed. Using a Grubbs’ test, we identify that the number of uniquely identified proteins in this replicate is an outlier, and therefore the protein abundances were justifiably neglected when calculating mean and standard deviations in the Mid CO_2_ treatment. Additionally, we calculate estimated enzyme specific activities by partitioning bulk CA activity for each treatment among the relative abundances of the two most abundant proteins, PtCA1 and Pt-ιCA. A detailed explanation of these calculations can be found in the Supporting Information Text S4 and Table S2.

## Results

### Growth rates

In the CO_2_ experiment, growth rates in the High CO_2_ treatment were significantly lower than both the Mid (*p* = 0.00009) and Low CO_2_ (*p* = 0.01) treatments (**Fig. 1a**, blue data points). In the Zn experiment, there was a significant positive growth effect as Zn^2+^ increased from 0.3 pM (Low Zn) to 3 pM (Mid Zn; *p* = 0.00004); however, as Zn^2+^ further increased to 10 pM (High Zn), no additional growth effect was observed (**Fig. 1a**, pink data points). For full growth curves, see Supporting Information Fig. S5.

### Carbonic anhydrase activity and protein abundances

Eleven unique CA proteins were identified in the proteome from these experiments (**Fig. 2a**; Supporting Information Fig. S6). PtCA1 and Pt-ιCA at Low Zn contributed the most substantially to the total CA protein abundance and showed clear responses to treatments (**Fig. 2b**). PtCA1 is a Zn-containing CA that localizes to the pyrenoid (Satoh et al. 2001; Tachibana et al. 2011) and its abundance decreases with pCO_2_ and increases with [Zn^2+^]. Pt-ιCA was initially named the *Low CO_2_ Inducible Protein of 63 kDa* by (Jensen et al. 2019); the authors additionally found this protein to localize to the chloroplast and exhibit a higher activity with Mn^2+^ in its active site relative to Zn^2+^, Ca^2+^, Co^2+^, or Mg^2+^. A new study believes to have identified the *ι*-CA operates without a metal cofactor under low Zn and pCO_2_ conditions in *P. tricornutum* (Avilan et al. 2026). Though this protein was originally named due to its production under low CO_2_, the concentrations of CO_2_(aq) in the seawater of the Low Zn and Low CO_2_ treatments were equivalent (Supporting Information Table S3), and its abundance increased only under conditions of Zn scarcity. We cannot claim whether it is operating with or without Mn in its active site and therefore continue to refer to it as Pt-ιCA. A list of the relative abundances of PtCA1 and Pt-ιCA in the six treatments can be found in Supporting Information Table S4. Additionally, the localizations of the remaining nine CAs (if known) are listed in Supporting Information Table S5.

CA activity showed strong treatment responses: it decreased with pCO_2_ in the CO_2_ experiments and increased with metal availability in the Zn experiments (Supporting Information Fig. S7). The Mid and High CO_2_ treatments had a lower CA activity relative to all other treatments. A decrease in CA activity between cultures grown with and without aeration has been previously documented (John-McKay and Colman 1997). CA activity exhibited an inverse relationship with concentrations of CO_2_(aq) in seawater at the time cells were harvested (**Fig. 2c**) and a direct relationship with the protein abundance of PtCA1 (**Fig. 2d**). Interestingly, though two different experimental conditions were carried out (bubbled vs. non-bubbled media), all points in the relationship between enzyme activity and [CO_2_]_aq_ collapse onto one line (**Fig. 2c**). Given the enhanced catalytic efficiency of Zn-containing CAs (Kim et al. 2020; Xu et al. 2008), the dominance PtCA1 exerts over the total CA protein abundance (**Fig. 2a**), and that its abundance corresponded linearly with CA activity (**Fig. 2d**), we hypothesize that majority of CA activity is attributable to PtCA1, followed by Pt-ιCA at low Zn concentrations. Using the co-measurement of bulk CA activity and individual CA proteins to create an over-determined system, we estimate the catalytic efficiency (estimated specific activity) of these two CAs from this experiment with some simplifying assumptions (Supporting Information Text S4). The operationally defined specific activity of PtCA1 was calculated to be 7 x 10^-14^ (activity peak intensity^-1^ sec^-1^) and Pt-ιCA to be 7 x 10^-16^ (activity peak intensity^-1^ sec^-1^).

### Active transport of HCO_3_^-^ via solute carrier proteins

Five unique bicarbonate transport proteins were identified in the proteomes from these experiments (Supporting Information Fig. S8). Of these, four are identified as part of the solute carrier 4 (SLC4) family HCO_3_^−^ transporter (Nakajima et al. 2013). SLC4-3 was abundant in all Zn treatments and its abundance decreased as pCO_2_ increased (**Fig. 3a**; pink bars). SLC4-4 was significantly more abundant in the High Zn treatment relative to both the Mid (*p* = 0.00006) and Low (*p* = 0.00045) Zn treatments and was scarcely present across treatments in the CO_2_ experiment (**Fig. 3a**; blue bars).

**Fig. 3.**
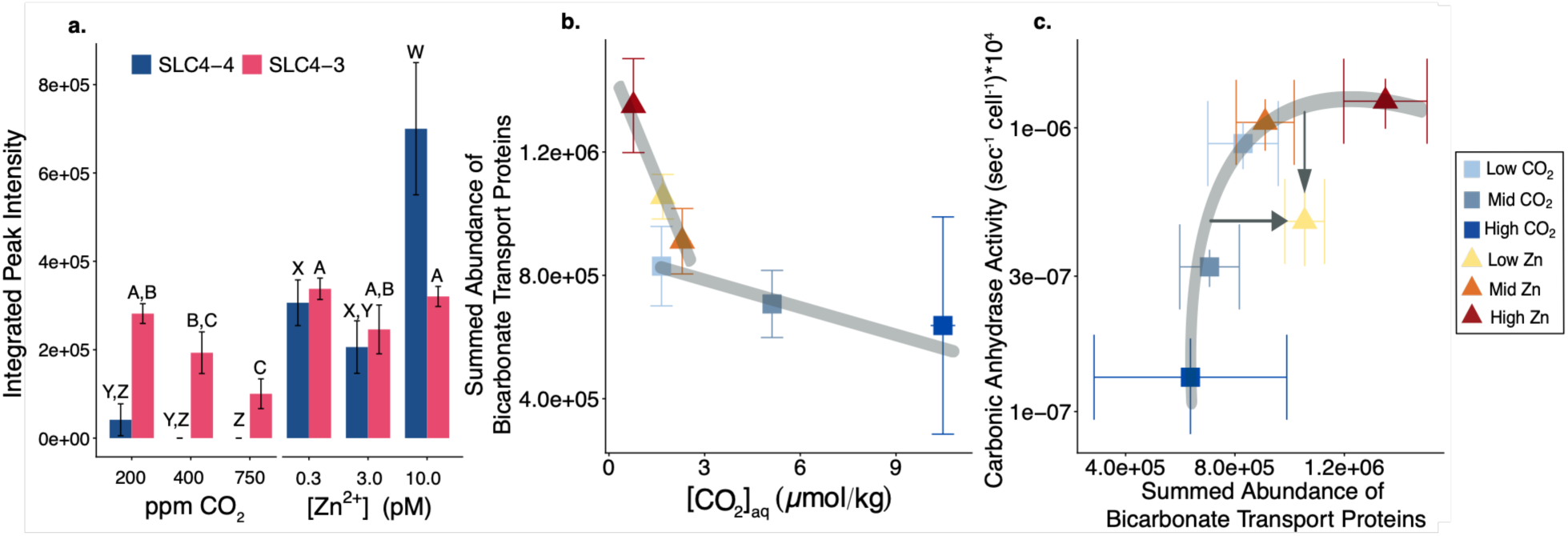
Bicarbonate transport protein abundance response to treatments. (**a**) Protein abundance of SLC4-3 (pink bars) and SLC4-4 (blue bars). Treatments with differing letters (A/B/C for SLC4-3; X/Y/Z for SLC4-4) reflect a significant difference in protein abundance from one another (p < 0.05; ANOVA followed by Tukey post hoc test). (**b**) The summed abundance of bicarbonate transport proteins plotted against the concentration of CO_2_(aq) in the seawater. Lines are hand-drawn to illustrated differences in slopes between the Zn and CO_2_ experiments. (**c**) CA activity plotted against the summed abundance of all bicarbonate transport proteins. Lines and arrows are hand-drawn to illustrate the deficit of CA activity relative to abundance of bicarbonate transport proteins at Low Zn. Data are reported as the average and standard deviation of biological triplicate.

The summed abundance of all bicarbonate transport proteins demonstrated an inverse relationship with the concentration of CO_2_(aq) in the seawater (**Fig. 3b**). The bubbled treatments in the CO_2_ experiment had a shallower slope relative to the non-bubbled treatments in the Zn experiment, consistent with a greater need for HCO_3_^-^ uptake without active CO_2_ supply. CA activity and active transport of bicarbonate appear to work in tandem with one another as exhibited by the linear relationship between the two parameters (**Fig. 3c**). There was an observed plateau of CA activity at approximately 10^-6^ sec^-1^cell^-1^ that met with a continued increase in bicarbonate transporter abundance in the non-bubbled Mid and High Zn treatments. Additionally, there was an excess of bicarbonate transport protein abundance in the Low Zn treatment relative to its measured CA activity. By performing a linear regression on the two variables in log-log space, we can compare the measured values of the total bicarbonate transport protein abundance and CA activity in the Low Zn treatment to their expected values as calculated from the equation (regression parameters are listed in Supporting Information Table S6). We find the measured CA activity is 1.7-fold lower than expected and the measured bicarbonate transport protein abundance is 1.3-fold higher than expected.

### Metal limitation in cultures

The presence and abundance of three key proteins (ISIP2A, ZCRP-A, and ZCRP-B) were used as indicators of metal limitation (Fe and Zn, respectively). ZCRP-A and ZCRP-B are two recently identified Zn responsive proteins in diatoms whose production are induced under conditions of Zn limitation (Kellogg et al. 2022). Both protein abundances were scarce in all but the Low Zn treatment (**Fig. 4a**).

**Fig. 4.**
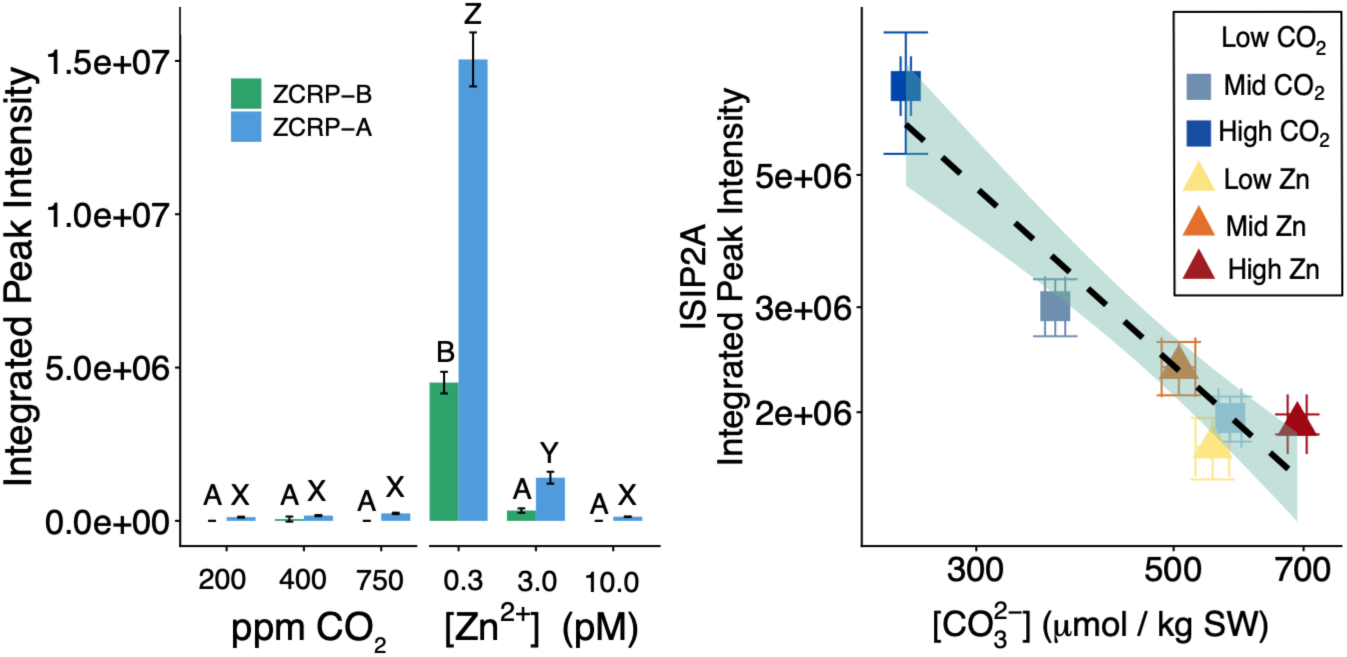
Demonstration of metal limitation in treatments. (**a**) Protein abundances of ZCRP-A (blue bars) and ZCRP-B (green bars). Treatments with differing letters (X/Y/Z for ZCRP-A; A/B for ZCRP-B) reflect a significant difference in protein abundance from one another (*p* < 0.05; ANOVA followed by Tukey post hoc test). (**b**) Protein abundance of the phytotransferrin ISIP2A versus carbonate ion concentration in seawater. Dashed line represents a linear regression model with an 80% confidence band in light green. Data are reported as the average and standard deviation of biological triplicate. For further information regarding regression parameters, refer to Supporting Information Table S6.

ISIP2A is a phytotransferrin that specifically utilizes FeCO_3_ complexes as a mechanism for iron uptake, and its responsiveness to iron scarcity has led to its use as a biomarker for cellular Fe stress (McQuaid et al. 2018). This protein was identified in all six treatments, though its abundance was significantly increased at High CO_2_ relative to all other CO_2_ and Zn treatments (*p* < 0.05 for all; **Fig. 4b**). Moreover, we observed a significant inverse relationship between its abundance and [CO_3_^2-^] (slope *p* = 0.029).

## Discussion

*P. tricornutum* is capable of high growth rates, resulting in high cell densities and depletion of the DIC pool, creating a potential for CO_2_ limitation. Due to the kinetic limitation of CO_2_ hydration, the consumption of CO_2_(aq) by photosynthesis is potentially faster than the rate at which DIC pool equilibrates (Riebesell et al. 1993). In large diatoms such as *Coscinodiscus granii* and *Ditylum brightwellii,* the production of extracellular CA mitigates this disequilibrium, as eCA interacts directly with the seawater in the DBL to facilitate a diffusive flux of CO_2_ into the cell, playing an essential role in supporting high rates of carbon fixation (Keys et al. 2026; W. Li et al. 2020). In contrast, *P. tricornutum* lacks significant eCA activity, likely due to its small size (20 μm long, 3 μm wide, Supporting Information Fig. S9) and its correspondingly low sensitivity to diffusive limitation. Therefore, one might presume its growth is highly sensitive to decreasing [CO_2_]_aq_, as its CCM function relies on uncatalyzed CO_2_ diffusion, iCA activity, and HCO_3_^-^ uptake (Burkhardt et al. 2001; Tachibana et al. 2011). Yet in this study, only a slight negative growth effect was observed as pCO_2_ decreased from 400 to 200 ppm, though this was met with a significant increase in CA activity (*p* = 0.013) implying a more efficient CCM at low pCO_2_ (**Fig. 1, 2c**). Furthermore, we observed no relationship between growth rates and CA activity (Supporting Information Fig. S10), drawing further contrast between diatoms that lack and express eCA (W. Li et al. 2020). The internal plasticity of *P. tricornutum* to respond to changing environmental conditions demonstrates the remarkable ability of its CAs (in conjunction with dynamic bicarbonate transporters, see below) to forestall CO_2_ limitation and maintain high growth rates given sufficient Zn.

### Zn availability influence on CA protein abundance and activity

In this study, though eleven different CA proteins were detected in the proteomes, the abundances of all but two (PtCA1 and Pt-ιCA) were relatively minor and did not show much variability among treatments (**Fig. 2a,b**). The presence of multiple CA proteins in *P. tricornutum* points to a degree of functional redundancy within the cell. That the cell is able to regulate its abundance of two CA proteins with resulting changes in overall activity demonstrates its sensitivity to metal, specifically Zn^2+^, availability. It is possible these CAs have roles outside photosynthesis and CCMs which have not yet been characterized and therefore would be regulated differently. Demetallized, or ‘apo’ enzymes are also possible, due to insufficient metal availability, which would contribute to the lack of differentiation in total CA protein abundances but large differences in overall activity.

In order for CA enzymes to exhibit catalytic activity, there must be sufficient metal availability for use as cofactors, and cellular metal allocation is assumed to be a highly regulated process (Sunda 1989). As an adaptive response to Zn scarcity, multiple isoforms of an enzyme can co-occur with different metal preferences, such as Pt-ιCA, observed to have more than 10-fold higher activity with a Mn cofactor than Zn (Jensen et al. 2019; Saito et al. 2008). Although, compared to other CAs (such as the β-CAs that use Zn cofactors), its activity is predicted to be much lower (Krishnamurthy et al. 2008; Xu et al. 2008). The preference for Zn as a cofactor for CA is likely due to its unique chemical properties: in brief, a Zn-bound hydroxide promotes rapid ligand exchange through a nucleophilic attack that catalyzes the CO_2_ hydration reaction (Kellogg 2022; Kim et al. 2020). In the current study, as calculated using paired abundance-activity measurements (Supporting Information Text S4, Table S2), the estimated specific activity of PtCA1 was 7 x 10^-14^ (activity peak intensity^-1^ sec^-1^) and Pt-ιCA was 7 x 10^-16^ (activity peak intensity^-1^ sec^-1^). These estimated activity measurements provide some basis for understanding the ecology of CA diversity, where Pt-ιCA is only synthesized under Zn scarcity duress, providing an alternate, albeit less efficient CA per enzyme unit. Specific activity measurements have been made for PtCA1 (Hopkinson et al. 2011) and Pt-ιCA (Jensen et al. 2019), but due to differences in methodology, the resulting kinetic parameters are not intercomparable.

The ability for *P. tricornutum* to increase its production of Pt-ιCA and instead use this ∼100-fold less efficient enzyme to retain partial CCM function demonstrates its biochemical flexibility when faced with Zn stress. Zn limitation inhibits many aspects of cell physiology, however, in the context of this manuscript, we attribute the observed decrease in growth rate at 0.3 pM Zn^2+^ as the outcome of a regulatory shift away from the efficient PtCA1 enzyme, lessening the resultant bulk CA activity and negatively impacting the CCM as well as other cellular Zn uses. Cultures grown with replete Zn can accomplish substantial CO_2_ drawdown from rapid growth rates, supported by a corresponding high CA activity. This is exhibited by the Low Zn and Low CO_2_ treatments which have equivalent concentrations of CO_2_(aq) but different proportions of PtCA1 and Pt-ιCA (Supporting Information Table S3, S4).That our estimated ESA_Pt-ιCA_ is two orders of magnitude lower than ESA_PtCA1_ justifies their difference in bulk CA activity. Moreover, the Low Zn treatment point is consistent with the line of regression between activity and PtCA1 protein abundance (**Fig. 2d**). By our calculations, if Pt-ιCA had a specific activity equal to that of PtCA1, this treatment would have a bulk CA activity much greater than what was observed, exceeding the linear relationship.

The *P. tricornutum* genome encodes for two Zn stress response proteins, which function as high-affinity ligands for metal uptake (ZCRP-B) and chaperones to the endoplasmic reticulum for use in protein synthesis (ZCRP-A). The concurrent increases in Pt-ιCA (**Fig. 2b**) and both ZCRP proteins (**Fig. 4a**) at 0.3 pM Zn^2+^ point to their role in maintaining homeostasis under conditions of Zn limitation. We posit the cell is allocating the limited Zn away from the CCM and towards protein biosynthesis, compensating with increased production of Pt-ιCA for partial restoration of the CCM.

It is well known that Cd and Co can substitute for Zn in CA either cambialistically or in different isoforms (Price and Morel 1990), however, their concentrations in the surface ocean can be 1-2 orders of magnitude lower than that of Zn and Mn (Kremling and Streu 2001). A potential Mn-based CA highlights a new and understudied role for this metal in the global carbon cycle. Mn is an essential micronutrient in algal photosynthesis due to its use in photosystem 2 and as a cofactor in various metalloenzymes such as superoxide dismutase, yet its cellular quota is estimated to be slightly lower than that of Zn (Twining and Baines 2013). Significant primary Mn limitation is difficult to demonstrate in cultures when concentrations of other trace metals such as Zn, Co, and Fe are replete (Brand et al. 1983; Sunda 1989; Sunda and Huntsman 1983). However, incubation experiments have shown stimulation of phytoplankton growth upon Mn addition in the Southern Ocean (Browning et al. 2021). The possibility for an isoform of marine CA to use Mn as a cofactor (or to function metal-free, as suggested by Avilan et al. (2026)) to retain partial function of their CCM demonstrates an advantageous biochemical flexibility and allows for Zn allocation to other metabolic processes such as protein synthesis.

### Fe stress response from High CO_2_

In addition to Zn and CO_2_ availability, growth rates are also affected by Fe availability: processes such as chlorophyll production and photosystem II efficiency become suppressed under Fe-limiting conditions, thus reducing growth rates (Allen et al. 2008). In this study, a negative effect on growth rate was observed in the High CO_2_ treatment (**Fig. 1**), which contrasts findings of a slight positive growth response to increased pCO_2_ in *P. tricornutum* (e.g., Y. Li et al. 2014; Wu et al. 2010; Ye et al. 2022). The prevaling conceptual model outlines that as pCO_2_ increases, a subsequent downregulation of the CCM permits greater energy allocation to additional metabolic processes, stimulating growth (Mackey et al. 2015). Yet, interactions between CO_2_, trace metals, and seawater can prevent this synergy.

ISIP2A (Iron Starvation Induced Protein-A) is a transferrin protein that binds the Fe-carbonate ion (CO_3_^2-^) pair during uptake (Morrissey et al. 2015). In a seawater-based growth medium where the majority of Fe is bound by EDTA, acidification decreases the concentration and availability of inorganic Fe species (Fe’), thereby decreasing Fe uptake rates (Shi et al. 2010). Moreover, as pH decreases, the carbonate equilibrium shifts in favor of CO_2_(aq) and away from CO_3_^2-^. Although photosynthesis increases pH and [CO_3_^2-^] through the consumption of CO_2_, bubbling at 750 ppm CO_2_ (High CO_2_) counters this effect and pushes DIC speciation away from CO_3_^2-^ (Supporting Information Table S3). A study by McQuaid et al. (2018) showed that as CO_2_ concentrations increased from 50 to 5,000 ppm, the rate of transferrin-mediated Fe’ uptake decreased and the expression of ISIP2A was induced. Indeed, in these experiments the inverse relationship between [CO_3_^2-^] and ISIP2A is consistent with the FeCO_3_ transport mechanism (**Fig. 4b**). If Fe uptake via the FeCO_3_ ion pair is a primary mechanism for cellular Fe acqustion, decreased metal uptake can explain the observed negative growth effect at High CO_2_.

### DIC acquisition via bicarbonate uptake in *P. tricornutum*

Bicarbonate is the largest component of DIC, which cells utilize via uptake through transport channel proteins (Nakajima et al. 2013). Active uptake of HCO_3_^-^ via a Na^+^ gradient is an energy intensive process. The overall abundance of bicarbonate transport proteins identified in the proteome (Supporting information Fig. S8) and the responsiveness of SLC4-4 and SLC4-3 to treatments (**Fig. 3a**) imply that active HCO_3_^-^ uptake plays a critical role in the *P. tricornutum*’s CCM, and several studies have shown this increases under CO_2_ limitation (Burkhardt et al. 2001; Nakajima et al. 2013; Tortell et al. 1997). The higher sensitivity in the relationship between bicarbonate transport protein abundance and [CO_2_]_aq_ in the non-bubbled Zn experiment treatments contrasts the single relationship observed between CA activity and [CO_2_]_aq_, which was independent of whether the treatments were bubbled or not (**Fig. 3c**). *P. tricornutum*’s CA enzymes are all located intracellularly, and they operate on the total intracellular DIC (i.e., diffusion of CO_2_ and active uptake of HCO_3_^-^) and therefore are less sensitive to the external seawater environment.

The reaction kinetics of DIC equilibration dictate the hydration of CO_2_ as the rate limiting step (in contrast to the rapid dehydration of HCO_3_^-^; Johnson, 1982). As RuBisCO can only fix CO_2_, HCO_3_^-^ uptake and intracellular CA activity are likely a tightly coupled process (**Fig. 3d**). As exhibited by the leveling off of CA activity yet continued increase in the abundance of bicarbonate transport proteins, we posit that *P. tricornutum* is accomplishing the remainder of DIC uptake not met through CO_2_ diffusion via active HCO_3_^-^ uptake. Though there could be a number of possibilities that explain the leveling off of CA activity, we detail two (non-exclusive) factors that seem most plausible to contribute to the observations here. First, CA enzymes operate at equilibrium, with a limited capacity to which they can elevate the concentration of CO_2_ in their vicinity. Thus, greater activity doesn’t inherently translate to a higher [CO_2_]_(aq)_. Secondly, there is a physical limitation on the surface area of the chloroplast membrane, where PtCA1 and Pt-ιCA localize, which can only accommodate so many enzyme copies. In the relationship between CA activity and bicarbonate transport protein abundance, we additionally observe a disproportionality higher abundance of bicarbonate transport proteins relative to the amount of CA activity per cell in the low Zn treatment (**Fig. 4c**; demonstrated by gray arrows) again implying *P. tricornutum*’s acclimation response to Zn limitation is to switch to an alternative mechanism of DIC acquisition – in this case active bicarbonate uptake – as an attempt to maintain growth.

### Hybrid C flow in *P. tricornutum*

Localization of key responsive proteins provides information about their function and role in the overall intracellular C transport. SLC4-1, 2, and 4 localize to the plasma membrane (Nakajima et al. 2013; Nawaly et al. 2023), and SLC4-6 and SLC4-7 are predicted to localize to the chloroplast membrane (Kroth et al. 2008). Both PtCA1 and Pt-ιCA localize to the chloroplast, and PtCA1 specifically localizes to the pyrenoid (Jensen et al. 2019; Tachibana et al. 2011). Together, the two groups of proteins serve to move DIC (either in the form of HCO_3_^-^ or CO_2_) from the seawater to the chloroplast, across multiple membranes, where CA and RuBisCO proteins are concentrated.

The findings from this study illustrate *P. tricornutum* utilizes an extremely effective carbon acquisition system that relies on a combination of CO_2_ diffusion and HCO_3_^-^ uptake that responds actively to environmental conditions. This C uptake system is also dependent on the interplay between metal availability and carbonate system speciation. Throughout its growth period in batch culture, the CO_2_(aq) becomes almost entirely depleted (Supporting Information Table S3) and the DIC demand likely exceeds what is possible by diffusion alone, highlighting the importance of the HCO_3_^-^ uptake pathway under CO_2_-limiting conditions. The data in this manuscript allow us to interpret a cellular model of C flow in *P. tricornutum* (**Fig. 5**). Here, we posit DIC enters through the cell both by uncatalyzed diffusion of CO_2_ via a concentration gradient and by active uptake of HCO_3_^-^. A portion of the HCO_3_^-^ brought into the cytoplasm where it rapidly equilibrates with CO_2_ via deprotonation. CO_2_ molecules diffuse across the chloroplast membrane(s) via a concentration gradient established from the rapid consumption of CO_2_ by RuBisCO. The other portion of HCO_3_^-^ gets transported to the site of C-fixation by SLC4 proteins embedded within the chloroplast. There, it is converted to CO_2_ by PtCA1 or Pt-ιCA (in scenarios of Zn limitation) and immediately consumed by RuBisCO. The K_M_ for CO_2_ is around 40 µM and its concentrations are likely much higher in the pyrenoid than in the chloroplast (Young et al. 2016). We hypothesize *P. tricornutum* is fixing CO_2_ at a rate that would prevent any outward diffusion or leakage. However, if there weren’t a sufficient sink of CO_2_, chloroplast CAs could be functioning to store excess DIC as HCO_3_^-^ (Hopkinson 2014). The shift to a Zn limiting environment promotes cells to synthesize high-affinity Zn ligands that transport the metal to the endoplasmic reticulum while using a potential Mn-containing CA isoform for CCM activity. As the concentration of CO_2_(aq) in the seawater increases, there is a corresponding decrease in bicarbonate transport proteins and increase in iron-carbonate transport proteins. The responsiveness of these proteins to changes in metal and CO_2_ availability highlights their role in DIC acquisition and the underlying systems biology of carbon adaptability.

**Fig. 5.**
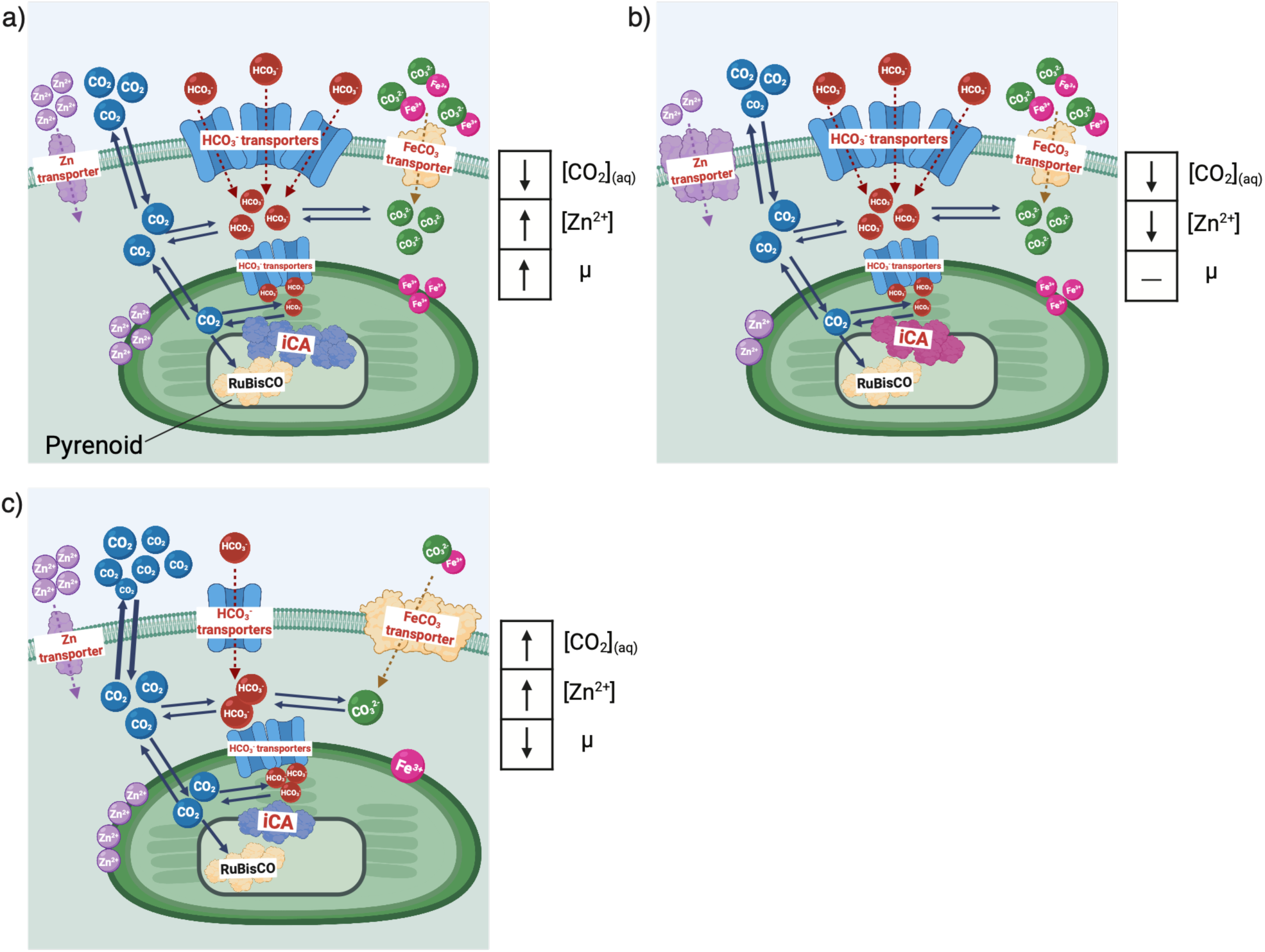
Schematic outlining carbon flow from the seawater diffusive boundary layer to chloroplast in a) low CO_2_ and high Zn, b) low CO_2_ and low Zn, and c) high CO_2_ and high Zn environments. Schematics were generated using stock images from BioRender.com without any use of AI.

## Acknowledgments

We thank Harriet Alexander and Sara Shapiro for use of and assistance with the Accuri flow cytometer, Loay Jabre for microscopy images of *P. tricornutum,* as well as all members of the Subhas and Saito labs. This work was funded by the National Science Foundation (OCE-2123055).

## Data Availability

The data that support the findings of this study are openly available at https://doi.org/10.26008/1912/bco-dmo.1010840.1 (physiology) and https://doi.org/10.26008/1912/bco-dmo.1011111.1 (protein abundances).

## Author Contributions

EAB: Conceptualization, Data curation, Formal analysis, Investigation, Methodology, Project administration, Resources, Software, Supervision, Validation, Visualization, Writing – original draft, Writing – review & editing

MAS: Funding acquisition, Conceptualization, Data curation, Formal analysis, Investigation, Methodology, Project administration, Resources, Supervision, Validation, Writing – review & editing

MGH: Data curation, Methodology

PZL: Data curation, Methodology

MRM: Data curation, Software

AVS: Funding acquisition, Conceptualization, Data curation, Formal analysis, Investigation, Methodology, Project administration, Resources, Software, Supervision, Validation, Writing – review & editing

