## Supporting Information Text for "Carbon concentrating mechanisms in a small coastal diatom demonstrate sensitivity to changes in Zinc and carbonate chemistry"

**This Supporting Information Contains the following:**

- 4 text sections
- 6 tables
- 10 figures

**Text S1: Carbonate chemistry measurements**

Carbonate chemistry samples were passed through a 0.2 µm filter (Whatman) to remove biomass and stored in 13 mL borosilicate vials where they were poisoned with mercuric chloride (0.02 % final concentration), sealed without headspace, and stored in the dark at room temperature prior to measurement. TA was measured using an open-system Gran titration on 1.5 mL samples brought to a total volume of 5 mL with Milli-Q water in technical duplicate using a Metrohm 805 Dosimat and a robotic Titrosampler (Borer et al., 2026). The precision of TA measurements assessed by the reproducibility of the standard was ± 2.5 µmol $kg^{-1}$. DIC was measured on 1.5 mL samples in technical duplicate using an Apollo SciTech AS-C6L DIC analyzer equipped with a LI-COR 7815 $CO_2$ detector (Wang et al., 2017). The precision of DIC measurements as calculated by the mean difference between technical duplicates was ± 4.2 µmol $kg^{-1}$. Measurements of TA and DIC were calibrated against in-house standards using seawater collected from Vineyard Sound, which were previously calibrated against seawater-certified reference materials (Batch #205) (Dickson, 2026). Measured values of DIC and TA were used as input parameters along with T, Sal, Total P, and Total Si in CO2SYS (Lewis et al., 1998) (ver. 2.3; using constants from Mehrbach et al. (1973) refit by Dickson and Millero (1987)) to calculate pH, $[HCO_3^-]$, $[CO_3^{2-}]$, and $[CO_2]_{aq}$.

**Text S2: Carbonic anhydrase activity analysis**

Carbonic anhydrase activity was assayed utilizing a membrane inlet mass spectrometer (MIMS), which measures ion intensities of dissolved gases based on detection of their mass to charge (m/z) ratio (Burlacot et al., 2020). A Pfeiffer QMG 220 mass spectrometer (closed-type ion source, 1–100 AMU mass range) was fitted with custom-made inlet consisting of a 6 inch long, 1/1600 O.D. stainless steel needle with 1 inch of perforations at the tip. Silastic tubing (0.23 mm thick) was gently slid over the needle to cover the perforations and was fixed by a knot at the tip of the needle. The needle functioned in tandem with the mass spec's high vacuum (operating at $4\times10^{-6}$ mbar) to pull dissolved gases from the sample or standard solution, without introduction of water, into the quadrupole mass spectrometer for detection.

Upon analysis, cell pellets were resuspended with 1 mL of a phosphate buffer medium (0.14 M sodium phosphate, pH 8). This slurry was transferred to a 15 mL centrifuge tube, where an additional 4 mL of buffer were added (Total volume = 5 mL) and the solution was sonicated (Microson Ultrasonic Cell Disruptor XL2000-001) in 3 bursts of 10 seconds at medium frequency, returning to ice intermittently. Through sonication, cells are lysed and any enzymatic activity would be in solution. The entire solution is assayed for activity following a 15 μL addition of spike. To prepare the spike solution, $^{13}C$-labeled sodium bicarbonate salt (85 mg; MilliporeSigma 870081-58-1) was equilibrated in 1 mL of $^{18}O$-labeled water (≥ 97% labeled $^{18}O$; MilliporeSigma 14314-42-2) for 24 hr. Samples were run in a conical reaction vial (MilliporeSigma Z115150) with a stir bar (MilliporeSigma 23227) gently agitating the solution and was held in a thermostat water jacket kept at 20°C.

All samples were assayed for their CA activity following a $^{18}O$-depletion method was followed, which tracks the ion currents of three different $^{13}C$ and $^{18}O$ labeled $CO_2$ isotopes over time as a double-labeled spike solution ($NaH^{13}C^{18}O_3$, m/z 49) interacts with the background buffer solution according to the following reaction;

$$^{13}C^{18}O_2 + H_2{}^{16}O \leftrightarrow H_2{}^{13}C^{16}O^{18}O^{18}O \leftrightarrow {}^{13}C^{18}O^{16}O + H_2{}^{16}O \leftrightarrow H_2{}^{13}C^{16}O^{16}O^{18}O \leftrightarrow {}^{13}C^{16}O^{16}O + H_2{}^{16}O \quad (\mathbf{1}).$$

The addition of the spike results in an initial increase in m/z 49, with subsequent increases in $^{13}C^{16}O^{16}O$ (m/z 45) and $^{13}C^{16}O^{18}O$ (m/z 47) as the spike reacts with the buffer's intense background $^{16}O$ signal. Utilizing the MIMS rapid response time, the interaction of the spike and the buffer are closely followed. The instrument takes approximately 130 measurements per minute, and each reaction is run for a total of 10 minutes. The fractional abundance of $^{18}O$ in $^{13}CO_2$ is used to calculate a rate of equilibration;

$$f^{18} = \frac{2*(49)+(47)}{2*(49+47+45)} \quad \textbf{(2)}.$$

Taking the slope of log(f18) versus time yields the rate of $CO_2$ hydration/dehydration (Subhas et al., 2019). This value is additionally multiplied by $10^4$ to get a whole number.

**Text S3: Proteomic analysis**

Harvested cell pellets were removed from the freezer and resuspended in lysis buffer (2% SDS, 50mM TEAB pH 8.5), and incubated at 95°C and mixed at 350 rmp for 10 (eppendorf; Thermomixer R). The samples were cooled on ice following the heated lysis buffer treatment, and 2 µL of benzonase nuclease (MilliporeSigma 707463) was added to each. This solution was left to incubate at 37°C for 30 min, shaking at 350 rpm. The extract was centrifuged at 14,100 × g for 10 minutes to remove cellular debris, and the extracted proteins in the supernatant were transferred to an ethanol washed tube. After the sample protein extraction, the total protein was quantified using a colorimetric assay at 562 nm (Pierce BCA total protein assay, Thermo Scientific 23225). Samples containing 25 µg of total protein were each subsampled into 200 µL of lysis buffer for the reduction, alkylation, and digestion of proteins.

To alkylate the samples, 4 µL of 500 mM dithiothreitol (DTT; Thermo Fisher Scientific 12-3-3483) in 50 mM ammonium bicarbonate (Ambic; Thermo Fisher Scientific 1066-33-7) was added and incubated at 45°C for 30 minutes. The alkylation was completed with an addition of 12 µL 500mM iodoacetamide (MilliporeSigma 144-48-9) in 50 mM Ambic, which incubated in the dark for 30 min at room temperature. Finally, the reaction was quenched with a second addition of 4 µL of DTT with a 10 min incubation and acidification using 12% phosphoric acid. Following extraction, reduction, alkylation, and acidification, the samples were transferred to an S-trap column (Protifi, S-Trap Mini Column) to clean, isolate, digest, and elute the proteins for analysis. Briefly, samples were thoroughly washed with binding buffer and a final wash with 90% methanol, digested with trypsin at a ratio of 25 µg protein:1 µg trypsin, left to incubate overnight at 37°C, and eluted with 50 mM Ambic, 50% formic acid, and 50% acetonitrile (ACN, Thermo Fisher Scientific A955-4) in 0.2% formic acid. This volume was quantified, and the purified protein was quantified using the BCA colorimetric assay. A protein concentration of 1 µg/µL was subsampled such that injection into the mass spectrometer would be uniform across all samples.

Tryptic peptides were analyzed with liquid chromatography coupled with tandem mass spectrometry (LS/MS/MS) where a Thermo Vanquish Neo HPLC reverse phase chromatography system was paired with a Thermo Scientific Fusion mass spectrometer using a Thermo Flex nanospray source. The peptides were eluted on a reverse phase C18 column (0.1 x 500 mm ID, 2 µm particle size, 120 Å pore size, C18 Reprosil-Gold, Dr. Maisch GmbH) using a 500 nl/min flow rate. All solvents were prepared with Fisher Optima grade reagents. Liquid chromatography was a nonlinear 60-minute gradient from 5%-95% buffer B, where A was 0.1% formic acid in water and B was 0.1% formic acid in ACN. Data independent acquisition (DIA) was used for analysis of the biological triplicates. The DIA scans comprised a precursor range of 380 m/z to 980 m/z on the Orbitrap with 24 m/z MS2 non-overlapping windows. The MS2 scans ranged from 150 to 2000 m/z, and the Orbitrap resolution was 60K for MS1 and 30K for MS2. The generated mass spectra from DIA analysis were searched against the translated genomes obtained from NCBI using MSFragger within FragPipe (v22.0: https://github.com/Nesvilab/FragPipe). An initial precursor mass tolerance of -10 to 10 ppm, a fragment mass tolerance of 10 ppm, a parent tolerance of 10.0 ppm, and a +57 carbamidomethyl fixed modification of cysteine were used, and MSBooster rescoring was used. Variable modifications were a +16 oxidation modification on methionine and a +42 acetyl modification on the N-terminus. Only peptides of length 6 to 50 and precursor charge states 2 to 4 were included, and a maximum of 1 missed cleavage was allowed. False discovery rates were calculated using decoys generated from a reverse database. A spectral library was generated from MSFragger, and DIA quantification was performed using DIA-NN to obtain a maximum protein false discovery rate of 0.01 (1%). An internal standardization within the DIA-NN software was performed based on MS1 signal intensity, such that reported integrated peak intensities between each sample were not over or underinflated (Demichev et al., 2020).

**Supporting Information Text S4: Estimated Enzyme Specific Activities of PtCA1 and Pt-ιCA**

PtCA1 and Pt-ιCA appear to be the two most treatment responsive proteins and here we estimate their specific activities. The total CA activity in a sample ($A_{tot}$) is the sum the of each enzyme's individual activity. For this exercise, $A_{tot}$ is partitioned three ways: PtCA1 and Pt-ιCA are considered independently (see Results and Discussion for justification; **Fig. 2**), and the remaining CA proteins are grouped into an 'other' term:

$A_{tot} = A_{PtCA1} + A_{Pt\text{-}\iota CA} + A_{other}$ **(1)**.

Enzyme specific activity (ESA) is traditionally defined as the enzyme's activity (A) per unit of protein. Using combined activity and proteomic measurements, we estimate an operationally defined ESA by partitioning $A_{tot}$ across the integrated peak areas for individual proteins, used as a weighting factor for protein abundance (Abu);

$A_{tot} = (ESA_{PtCA1} * Abu_{PtCA1}) + (ESA_{Pt\text{-}\iota CA} * Abu_{Pt\text{-}\iota CA}) + (ESA_{other} * Abu_{other})$ **(2)**.

For treatments in which the relative abundance of Pt-ιCA ($Abu_{Pt\text{-}\iota CA}$) is negligible (Supporting Information **Table S4**), $A_{Pt\text{-}\iota CA}$ could be ignored, thus **Equation 2** is reduced to the following:

$A_{tot} = (ESA_{PtCA1} * Abu_{PtCA1}) + A_{other}$ **(3)**.

A regression of Atot against $Abu_{PtCA1}$ for these treatments yielded values for $ESA_{PtCA1}$ (the slope) and $A_{other}$ (the intercept). For the treatment with significant Pt-ιCA protein abundance, **Equation 2** was then rearranged to solve for $ESA_{Pt\text{-}\iota CA}$:

$ESA_{Pt\text{-}\iota CA} = (A_{tot} - (ESA_{PtCA1} * Abu_{PtCA1}) - A_{other})/(Abu_{Pt\text{-}\iota CA})$ **(4)**.

For values used to calculate ESA, see Supporting Information Table S2. These calculations provide a conservative estimate for ESA, due to assumptions that must be made. Most significantly is the assumption that every enzyme has activity, as our methods cannot differentiate between enzymes with and without activity. Standard proteomic methods cannot differentiate active enzymes from metal free apo-enzymes, although metalloproteomic techniques are capable of detecting metalation (Saito & McIlvin, 2025).

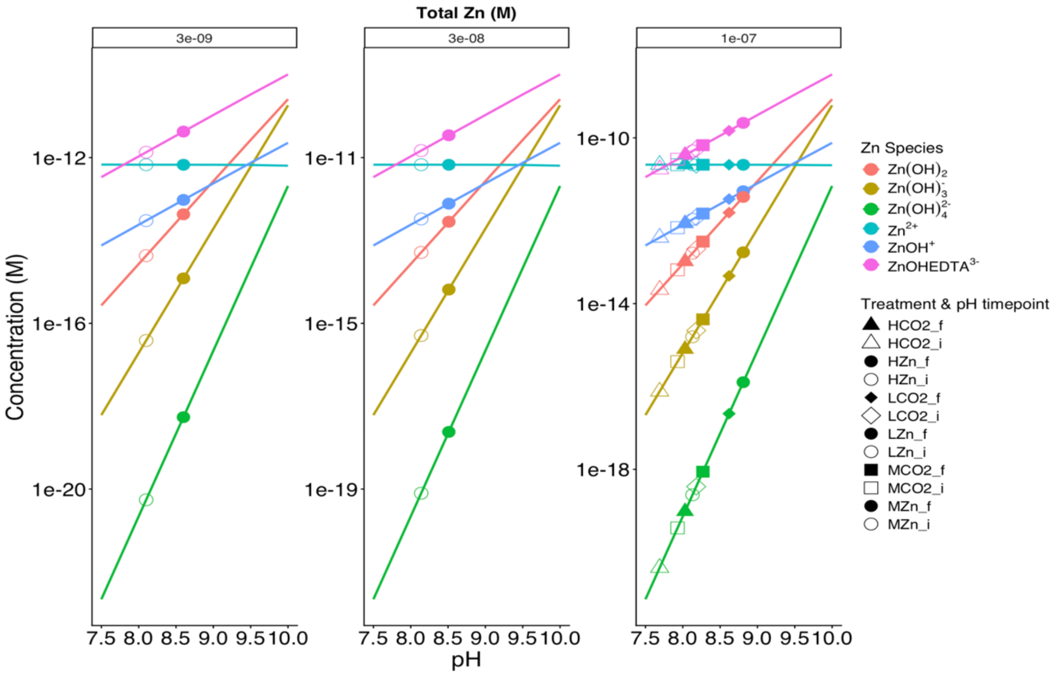


**Fig S1.** Concentration of selected inorganic and EDTA-bound Zn species versus pH at the three $[Zn]_T$. Colors denote the various inorganic species, with symbols representing the initial (open) and final (filled) pH values of experimental treatments. In addition to the treatment pH values, speciation calculations were done at pH 7, 9, 9.5, and 10, though symbols are omitted for clarity.

Zn speciation calculations were carried out with chemical equilibria software (MINEQL+ v. 5.0) using experimental data for comparison to the original calculations done by Sunda and Huntsman (1995). The concentrations of $Mg^{2+}$ (0.053 M), $Ca^{2+}$ (0.01 M), $SO_4^{2-}$ (0.0283 M), $PO_4^{3-}$ ($3.6 \times 10^{-5}$ M) as well as an ionic strength of 0.5 M were considered in these calculations.

Across 2.5 pH units there are large changes in the concentrations of various inorganic Zn species, though these species represent a small fraction (< ~2%) of the total Zn pool. At a constant $[Zn]_T$, there is a negligible change in $[Zn^{2+}]$ with pH due to EDTA. At 100 nM $Zn_T$, $[Zn^{2+}] = 2.25 \times 10^{-11}$ M, at 30 nM $Zn_T$, $[Zn^{2+}] = 6.7 \times 10^{-12}$ M, and at 3 nM $Zn_T$, $[Zn^{2+}] = 6.7 \times 10^{-13}$ M.

This system is in contrast to that of Fe, where there are large changes in inorganic Fe uptake rates with pH even in an EDTA buffered system (Shi et al., 2010). This distinction is due to the strong complexation by Fe hydroxides, which comprises a much greater fraction of the total Fe.

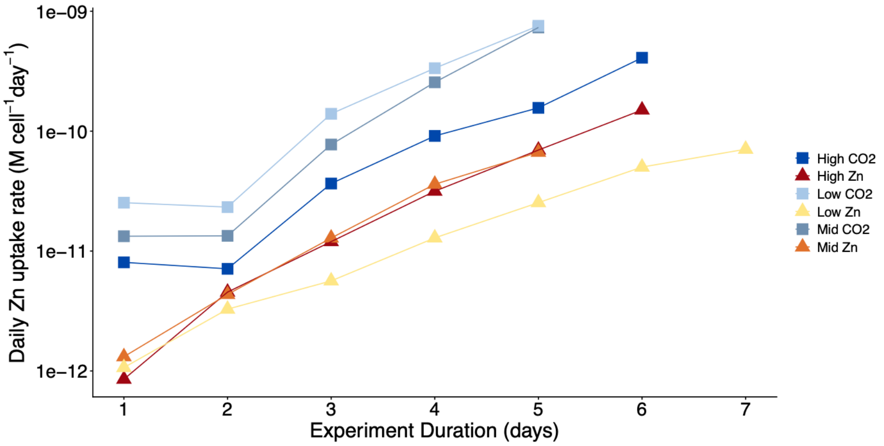


**Fig S2.** Estimated daily Zn uptake rates across the 6 experimental treatments, using a single Zn quota value. These calculations were done to verify the Zn inventory in the media was not depleted. At steady state (as observed by the linear growth curve; Sunda and Huntsman (1995)), Zn uptake can be estimated from the product between metal quota and growth rate using the following equation; $\rho = Q*\mu$. The Zn metal quota of 0.5 mmol Zn mol $P^{-1}$ for *P. tricornutum* was previously measured in Kellogg et al. (2020). 5.5E-15 mol P $cell^{-1}$ was used to convert the Zn metal quota to a per cell basis. Instantaneous growth rates ($day^{-1}$) and daily cell concentration (cells $mL^{-1}$) were used to convert the uptake rate to units of M $day^{-1}$.

**Table S1.** Percentage of $Zn^{2+}$ taken up relative the total added Zn to the media at the final timepoint of each experimental treatment.

| | |
|---|---|
| High Zn | 0.15 % |
| Mid Zn | 0.22 % |
| Low Zn | 2.36 % |
| High $CO_2$ | 0.41 % |
| Mid $CO_2$ | 0.73 % |
| Low $CO_2$ | 0.76 % |

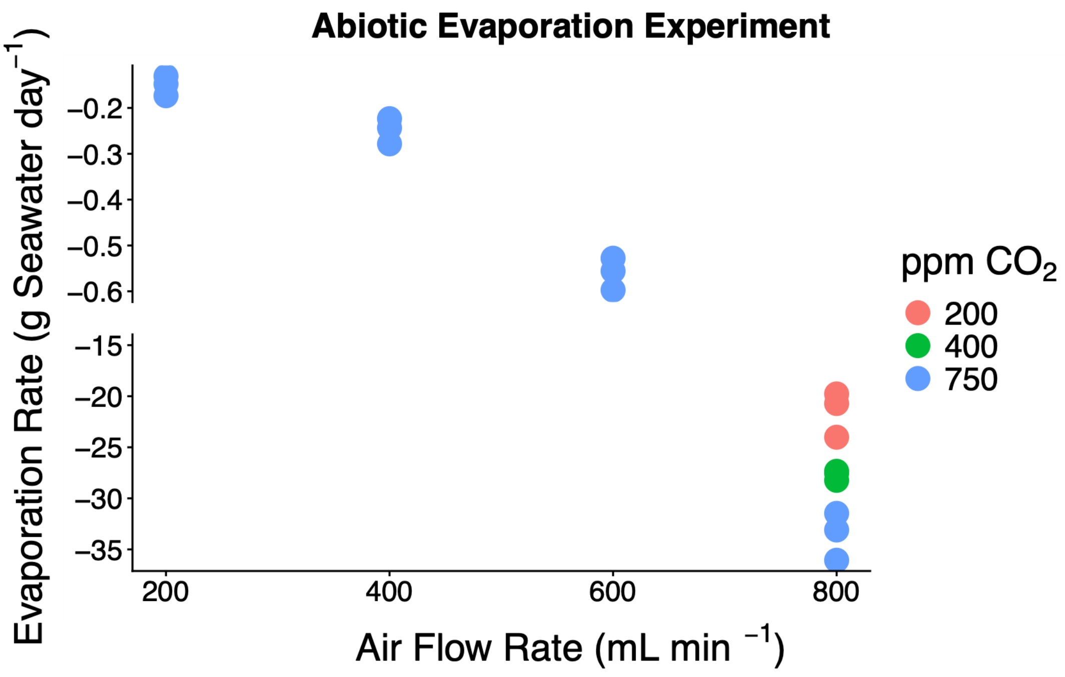


**Fig S3**. Evaporation rate of Milli-Q water measured as mass loss over time at a given air flow rate colored by $pCO_2$ treatment. In the ebullition design, a single air flow rate was maintained across all treatments. Flow rates of $CO_2$ were manipulated such that resulting gas streams had measurable $pCO_2$ values of 200, 400, and 750 ppm as determined by the binary mixing equation,

desired $pCO_2$ of treatment = $[(1\%\ CO_2) \times (Q_{CO2})] \div (Q_{CO2} + Q_{Air})$

where $Q$ is flow rate in mL min$^{-1}$. At higher air flow rates, there is more turbulent bubble injection, and higher evaporation rates. Evaporation of seawater skews DIC and TA concentrations higher and alters the environment in which the cultures are growing. At an air flow rate of 400 mL min$^{-1}$, evaporation was found to be less than 0.3 g SW day$^{-1}$, or 0.12% of the total volume per day.

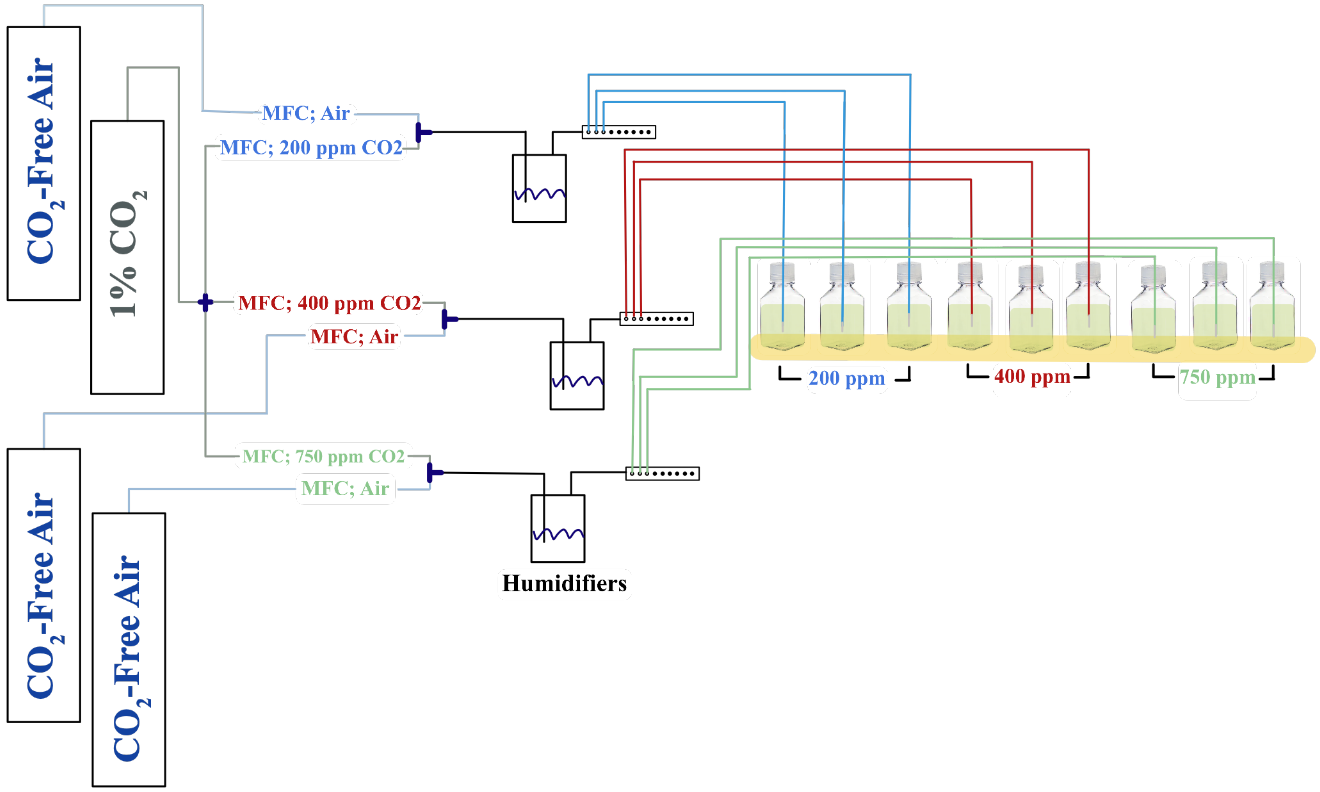


**Fig S4.** Schematic of gas mixing design for $CO_2$ experiments. Lines before humidifiers carry unmixed gas streams of $CO_2$-free air (blue) and 1% $CO_2$ (gray). The two gasses are mixed to reach the target $pCO_2$: 200 ppm (blue lines), 400 ppm (red), and 750 ppm (green). Length of lines are not to scale. All three $pCO_2$ treatments had independent pathways from the gas cylinder downstream to the culture bottles. One $CO_2$ tank (1% $CO_2$ in air) was split three ways using mass flow controllers (Tylan FC 260). Each $pCO_2$ treatment had an independent $CO_2$-free air tank with its flow rate regulated by a mass flow controller (Alicat Basis 2). The air and $CO_2$ gas streams were mixed and immediately humidified. Following humidification, each $pCO_2$ gas stream was split three ways using a manifold (long rectangle, where the open circles represent ports where bottle lines are connected, and filled circles represent ports which are blocked off) for biological triplicate measurements. The tubing is connected by a luer lock port drilled into the cap of the culture bottle. Prior to the gas stream entering the bottle, the end of each line was filtered (25 mm, 0.2 µm Whatman filter). The mean flow rate of gas to each bottle was 64.3 ± 1 mL $min^{-1}$. Inside the bottle, the end of the tubing is connected to an air diffuser to create smaller, more frequent bubbles. Finally, the culture bottles which sit above a strip of fluorescent light, as indicated by the yellow bar.

**Table S2.** Values used to calculate the Enzyme Specific Activities for PtCA1 and Pt-ιCA. Units for Activity are reactions per second per cell and protein abundance is measured as the integrated peak intensity.

| | |
|---|---|
| **Slope**; $ESA_{PtCA1}$ (Activity / Abundance) | 6.53E-14 |
| **Intercept**; $ESA_{other} * Abu_{other}$ (Activity) | 4.64E-08 |
| ${}^{LZn}MeasAct$ (Activity) | 4.68E-07 |
| ${}^{LZn}Abu_{PtCA1}$ (Abundance) | 6.32E+06 |
| ${}^{Lzn}Abu_{Pt-\iota CA}$ (Abundance) | 1.28E+07 |
| ${}^{Lzn}Abu_{other}$ (Abundance) | 5.88E+06 |
| $ESA_{Pt-\iota CA}$ (Activity / Abundance) | 1.28E-15 |

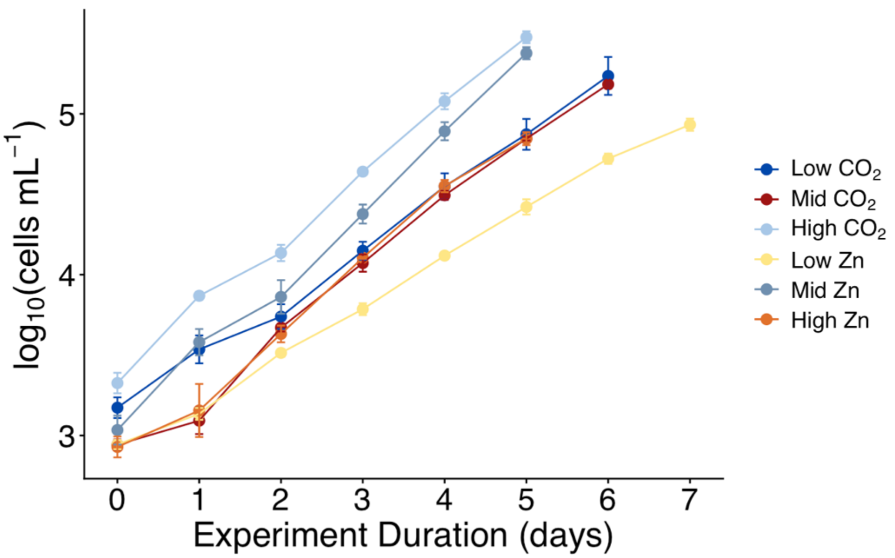


**Fig S5**. Growth curves plotted as cell density throughout experiment duration colored by treatment. Points represent the average cell density of biological triplicate with error bars denoting standard deviation.

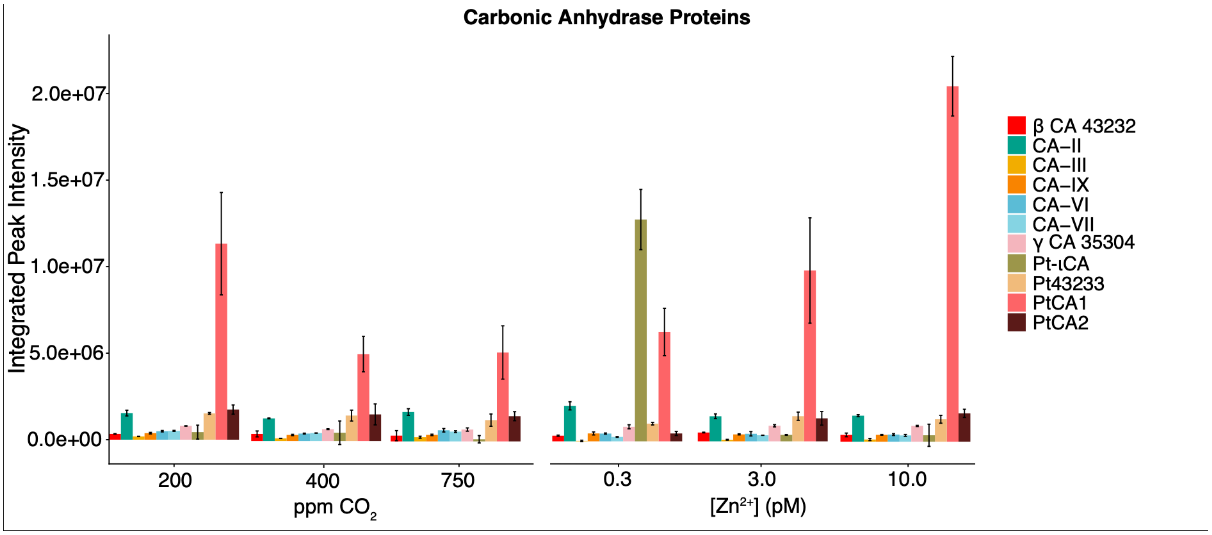


**Fig S6**. All carbonic anhydrase proteins and their abundances identified in the proteome of *P. tricornutum* within the $CO_2$ and Zn experiments. Bars depict the mean protein abundance of biological triplicate with error bars depicting standard deviation.

**Table S3.** Description of the six experimental treatments and carbonate chemistry values measured at the termination of the experiment. Measured values are reported as the average with error reported as standard deviation of biological triplicate.

| Exp. type | Treatment name | $[Zn^{2+}]$ (pM) | DIC (µmol $kg^{-1}$) | Alk (µmol $kg^{-1}$) | pH | $[HCO_3^-]$ (µmol $kg^{-1}$) | $[CO_3^{2-}]$ (µmol $kg^{-1}$) | $[CO_2]_{aq}$ (µmol $kg^{-1}$) |
|---|---|---|---|---|---|---|---|---|
| $CO_2$ | Low (200 ppm) $CO_2$ | 10.23 | 1727 ± 24 | 2578 ± 3 | 8.62 ± 0.03 | 1148 ± 41 | 539 ± 17 | 2.0 ± 0.2 |
| | Mid (400 ppm) $CO_2$ | 10.23 | 1985 ± 3 | 2518 ± 16 | 8.27 ± 0.01 | 1645 ± 8 | 334 ± 9 | 6.1 ± 0.2 |
| | High (750 ppm) $CO_2$ | 10.23 | 2162 ± 24 | 2515 ± 22 | 8.02 ± 0.01 | 1929 ± 26 | 220 ± 3 | 12.8 ± 0.5 |
| Zn | Low Zn | 3.07 | 1687 ± 34 | 2508 ± 4 | 8.60 ± 0.04 | 1172 ± 57 | 514 ± 23 | 2.0 ± 0.3 |
| | Mid Zn | 0.31 | 1775 ± 27 | 2520 ± 7 | 8.51 ± 0.04 | 1303 ± 48 | 469 ± 21 | 2.8 ± 0.3 |
| | High Zn | 10.23 | 1542 ± 19 | 2575 ± 9 | 8.81 ± 0.03 | 897 ± 35 | 644 ± 16 | 0.9 ± 0.1 |

**Table S4**. Proportion of PtCA1 and Pt-ιCA relative to the summed abundance of all CA proteins, expressed as percentage.

| Treatment | % PtCA1 Abundance ± Error | % Pt-ιCA Abundance ± Error |
|---|---|---|
| Low Zn | 25 ± 6 | 51 ± 8 |
| Mid Zn | 58 ± 21 | 2 ± 0.4 |
| High Zn | 74 ± 8 | 1 ± 2 |
| Low $CO_2$ | 59 ± 18 | 2 ± 2 |
| Mid $CO_2$ | 41 ± 10 | 4 ± 6 |
| High $CO_2$ | 41 ± 14 | 1 ± 2 |

**Table S5.** ProteinID, accession numbers, and localization (if applicable) of relevant proteins identified in the proteome of *P. tricornutum.* Proteins without listed localizations have yet to be identified to our knowledge. Proteins without references were identified when the genome was assembled and annotated in Bowler et al. (2008) and have not been studied further to our knowledge.

| **Protein Name** | **Subclass** | **Protein ID** | **Accession Number** | **Localization** | **Ref.** |
|---|---|---|---|---|---|
| CA-II | α | B7FUE7 | XP_002178308 | Chloroplast (periplastidal compartment) | Tachibana et al., (2011) |
| CA-III | α | B7GA80 | XP_002183900 | Plastid | Tachibana et al., (2011) |
| CA-IX | γ | B5Y401 | XP_002185753 | Mitochondria | Tachibana et al., (2011) |
| CA-VI | α | B7FUE9 | XP_002178310 | Chloroplast endoplasmic reticulum | Tachibana et al., (2011) followed by Samukawa et al., (2014) |
| CA-VII | α | B7FNT2 | XP_002176590 | Chloroplast endoplasmic reticulum | Tachibana et al., (2011) followed by Samukawa et al., (2014) |
| PtCA1 | β | B7FNU0 | XP_002176594 | Chloroplast (pyrenoid) | Kitao et al. (2008) |

| | | | | | |
|---|---|---|---|---|---|
| | | | | | followed by Tachibana et al., (2011) |
| CA 43232 | β | B7FR27 | XP_002177506 | | |
| Pt43233 | θ | B7FR28 | XP_002177507 | Thylakoid lumen penetrating the pyrenoid | Kikutani et al., (2016) |
| PtCA2 | β | B7FXP8 | XP_002179805 | Chloroplast | Kitao et al., (2008) followed by Tachibana et al., (2011) |
| CA 35304 | γ | B7FY50 | XP_002179677 | | |
| LCIP63 | I | B7G7G0 | XP_002183267 | Chloroplast | Jensen et al., (2019) |
| SLC4-4 | N/A | B7S436 | XP_002176330 | Plasma membrane | Nakajima et al., (2013) followed by Nawaly et al., (2023) |
| SLC4-3 | N/A | B7S4D4 | XP_002176377 | | Nakajima et al., (2013) |
| SLC4-6 | N/A | B7FQY4 | XP_002177487 | Predicted: chloroplast membrane | Kroth et al (2008) followed by Nakajima et al., (2013) |
| SLC4-7 | N/A | B7FYF1 | XP_002179925 | Predicted: chloroplast membrane | Kroth et al (2008) followed by Nakajima et al., (2013) |
| Putative $HCO_3^-$ transporter | N/A | B5Y5V6 | XP_002186254 | | |
| ISIP2A | N/A | B7FYL2 | XP_002179762 | Cell surface membrane | Morrissey et al (2015) |
| ZCRP-A | N/A | B7GEE7 | XP_002185475 | Chloroplast endoplasmic reticulum | Kellogg, Moosburner, et al.,(2022) |
| ZCRP-B | N/A | B7G9Q2 | XP_002183815 | Cell membrane | Kellogg, Moosburner, et al.,(2022) |

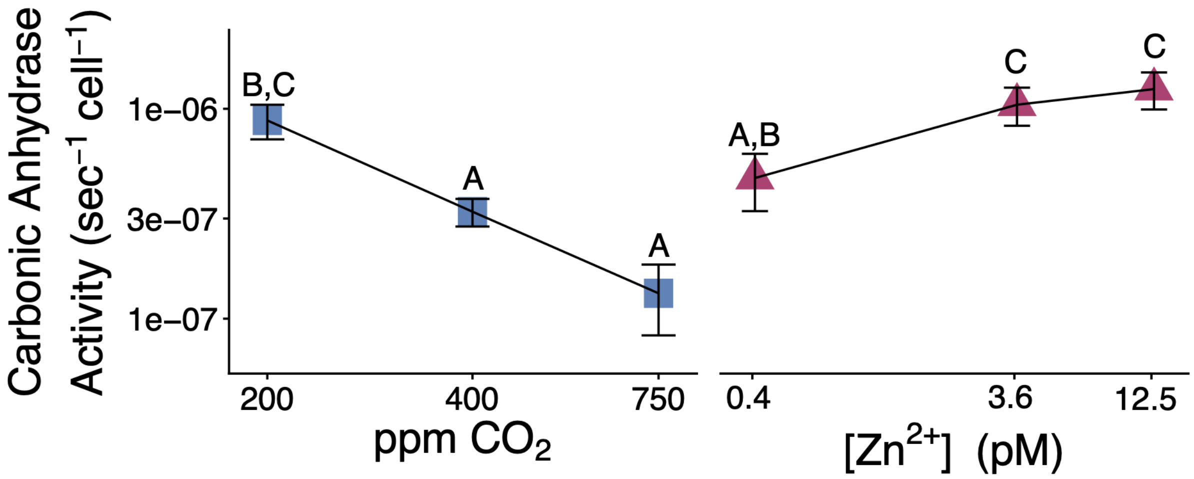


**Fig S7.** Carbonic anhydrase activity across the six treatments with pairwise comparison significance (ANOVA $p$ = 1.97E-7). Points represent the average of biological triplicate with error bars reported as standard deviation (n = 3). Treatments with matching letters reflect an insignificant difference in protein abundance from one another ($p < 0.05$; ANOVA followed by Tukey post hoc test).

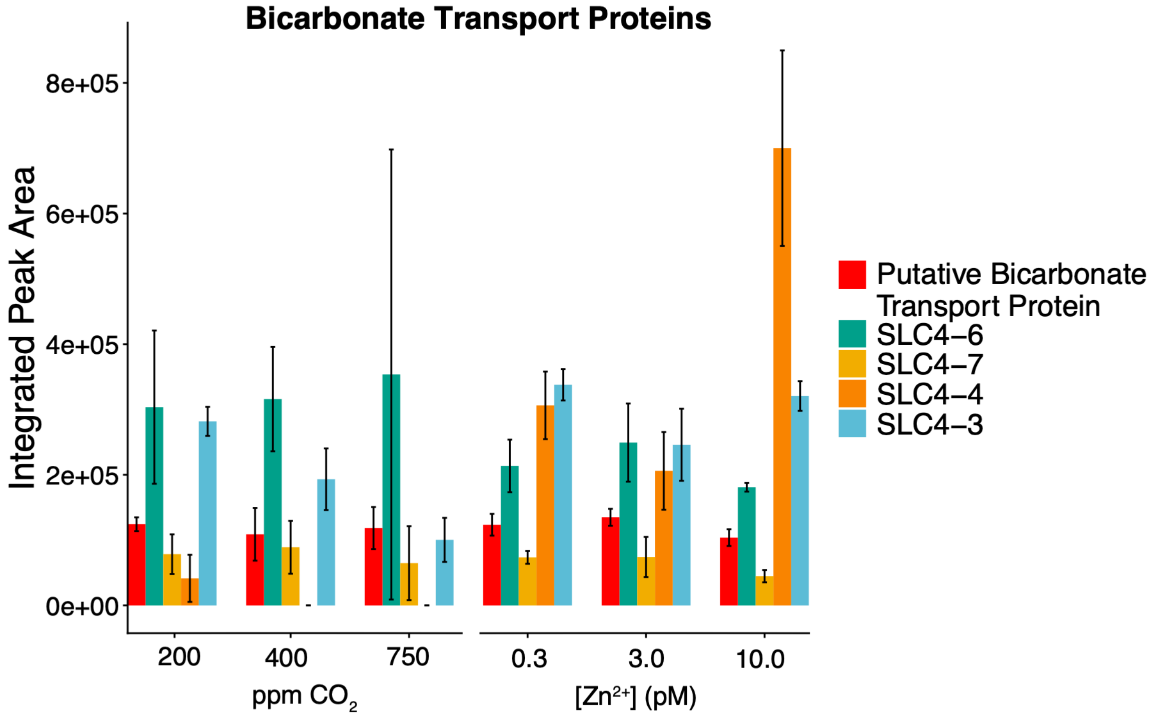


**Fig S8**. All bicarbonate transport proteins and their abundances identified in the proteome of *P. tricornutum* within the $CO_2$ and Zn experiments. Bars depict the average protein abundance of biological triplicate with error bars depicting standard deviation.

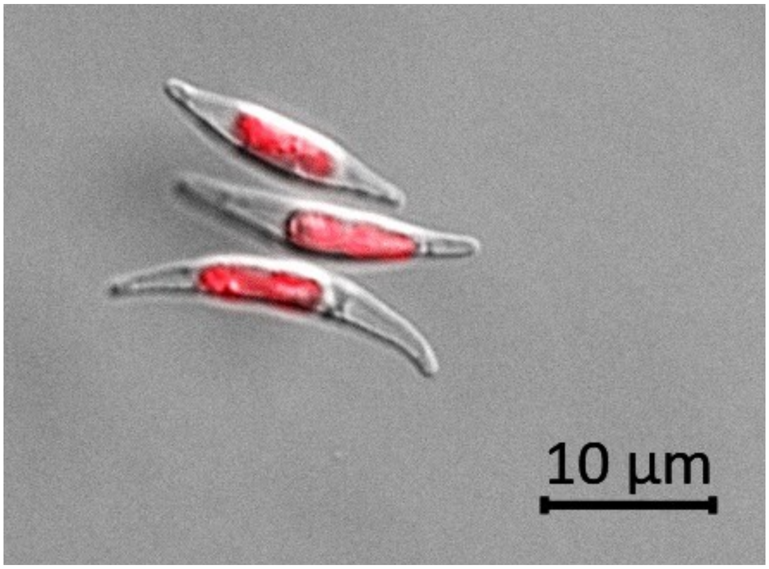


**Fig S9.** Microscopy image of *P. tricornutum*, taken by Loay Jabre.

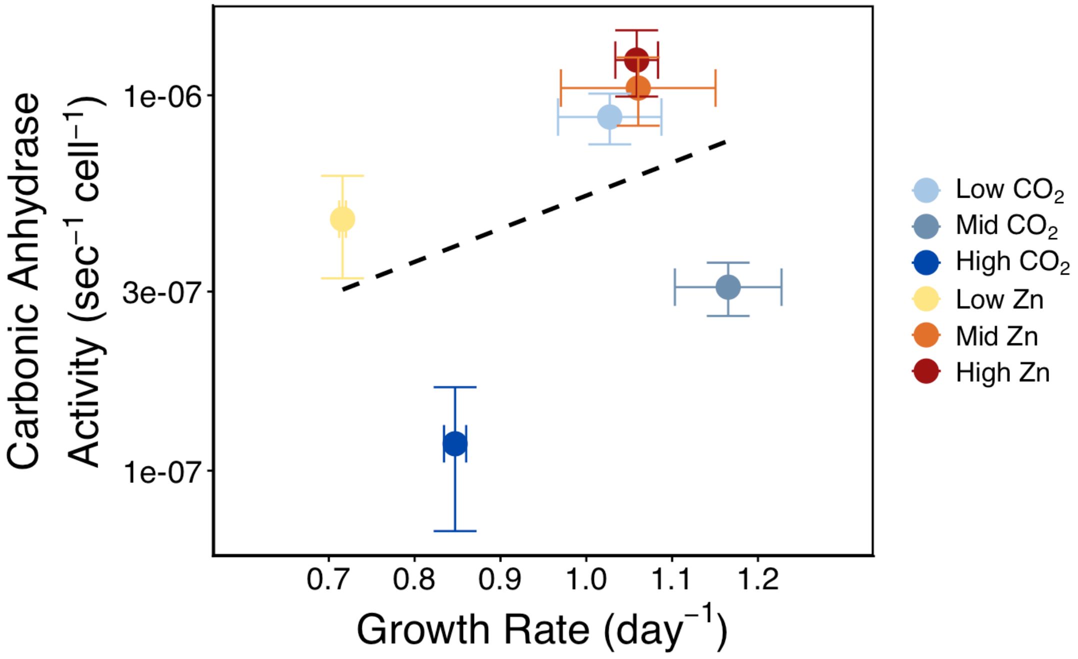


**Fig S10.** Carbonic anhydrase activity plotted against a sample's corresponding growth rate. Points represent average of biological triplicate with error bars reporting standard deviation. The slope of this line was 1.1e-06 ± 1e-06 ($p = 0.44$) and the intercept was -3.6e-07 ± 1e-06 ($p = 0.78$).

Growth rate is a direct measurement of the rate at which the cell is fixing C, assuming a constant cellular C quota across treatments. CA activity has a strong influence on the rate of delivery of $CO_2$ to the site of C-fixation in the chloroplast and could result in a positive correlation with growth rate. In these experiments, we observed a strong decoupling between CA activity and growth rate, potentially explained by the consistent delivery of $CO_2$ to the growth media achieved through bubbling. One example of this is the Mid $CO_2$ treatment, which exhibited the highest growth rate, but a lower CA activity relative to the Low $CO_2$, Mid and High Zn treatments. Another example is the High $CO_2$ treatment, which had a higher growth rate but lower CA activity than that of the Low Zn treatment.

**Table S6**. Parameters from linear regression fittings.

| **Fig #** | **X parameter** | **Y parameter** | **Slope ± Error** | **Intercept ± Error** | $\mathbf{R^2}$ |
|---|---|---|---|---|---|
| 5a | $[CO_2]$(aq) | Carbonic Anhydrase activity | -7.76-08 ± 3.0e-08 | 1.02e-06 ± 1.8e-07 | 0.6257 |
| 5b | PtCA1 Integrated Peak Intensity | Carbonic Anhydrase activity | 6.514e-14 ± 1.781e-14 | 4.951e-08 ± 1.976e-07 | 0.7699 |
| 6 | ISIP2A Integrated Peak Intensity | $[CO_3^{2-}]$ | -1.14e+04 ± 3.4e+04 | 8.19e+06 ± 1.6e+06 | 0.7365 |
| 7b | Sum of bicarbonate transport proteins Peak Intensity | Carbonic Anhydrase activity | 2.392 ± 1.003 | -20.503 ± 5.97 | 0.5872 |
